# On the importance of accounting for intraspecific genomic relatedness in multi-species studies

**DOI:** 10.1101/321927

**Authors:** Simon Joly, Dan F. B. Flynn, Elizabeth Wolkovich

## Abstract

1. Analyses in many fields of ecology are increasingly considering multiple species and multiple individuals per species. Premises of statistical tests are often violated with such datasets because of the non-independence of residuals due to phylogenetic relationships or intraspecific population structure. If comparative approaches that account for the phylogenetic relationships of species are well developed and their benefits demonstrated, the importance of considering the intraspecific genetic structure, especially in combination with the phylogenetic structure, has rarely been addressed.
2. We investigated whether it is beneficial to account for intraspecific genomic relatedness in multi-species studies. For this, we used a Phylogenetic Mixed Model to analyze first a suite of simulated data and then results from one example ecological study—a budburst experiment where clippings of 10 tree and shrub species were subjected to different temperatures and photoperiods.
3. We found that accounting for intraspecific genetic structure yields more accurate and precise fixed effects as well as increased statistical power, but more so when the relative importance of the intraspecific to the phylogenetic genetic structure is greater. Analysis of the budburst experiment further showed that accounting for intraspecific and phylogenetic structures yields improved estimates of warming and photoperiod effects and their interaction in explaining the time to budburst.
4. Our results show that statistical gains can be made by incorporating information on the intraspecific genomic relatedness of individuals in multi-species studies. This is relevant for investigations that are interested in intraspecific variation and that plan to include such observations in statistical tests.

## Introduction

The reactions of different species to external stimuli are not independent. Because physiological responses have a genetic basis, closely related species are more likely to have similar responses to a specific treatment. This phylogenetic non-independence of species responses violates the assumptions of most statistical tests, such as the independence of residuals in regression, and negatively impacts the results in terms of parameter estimates and p-values (e.g., Revell, 2010). This has been recognized for some time and a family of methods—comparative methods—have been developed to address this problem (Felsenstein, 1985; Grafen, 1989; Lynch, 1991; Ives et al., 2007; Felsenstein, 2008; Revell, 2010; Hadfield and Nakagawa, 2010).

Recently there has been growing awareness among ecologists of the need to also consider intraspecific variation within ecophylogenetic analyses. Within community ecology, there have been calls to increase studies of intraspecific trait variation (Violle et al., 2012; Alofs, 2016) and to develop the necessary statistical models for such multilevel data (Funk et al., 2017; Read et al., 2016). Similarly, studies of climate change have repeatedly highlighted the need for models that incorporate variation in responses across both species and populations (Willis et al., 2008; Charmantier et al., 2008; Anderson et al., 2009; Chen et al., 2011). However, little attention has been given to the genetic correlation structure present below the species level within the field of comparative methods (but see Hansen et al., 2000; Felsenstein, 2002; Stone et al., 2011; Read et al., 2016; Garamszegi, 2014). Moreover, studies rarely account for both phylogenetic and intraspecific genetic correlations simultaneously, even though the sampling structure in many ecological studies calls for such a design.

Presently, studies that account for phylogenetic correlation almost always ignore intraspecific genetic structure and as such assume that intraspecific samples are drawn from a single population. In contrast, many ecological studies explicitly sample individuals across important geographical ranges or from populations among which gene flow could be restricted, resulting in a potentially non-trivial correlation structure among samples. If this correlation is important, statistical tests that do not account for it are expected to be biased.

Until recently, the difficulty of obtaining genetic data to accurately estimate intraspecific genetic correlations provided sufficient justification for ignoring this source of variance in ecological studies. But the rapid development of sequencing techniques now allows precise estimation of genetic relatedness (Gienapp et al., 2017), at relatively affordable prices. This could allow a better understanding of how ecological responses are influenced by the genetic relationships among species and populations at once. One area where this potential is particularly high is climate change research, where evidence of rapid ecological and evolutionary change is growing. Research has highlighted that species responses to climate change appear phylogenetically patterned, with species from certain clades and with particular traits appearing most vulnerable to local extinctions with warming (Willis et al., 2008). At the same time other work has highlighted discrepancies in species responses when studied over space (Charmantier et al., 2008), suggesting populations within species may show different responses to climate change. This is supported by population-level research that has found large differences in the range shifts of northern versus southern populations with warming (Anderson et al., 2009; Chen et al., 2011). Such results make clear that the best estimates of responses will require methods that consider variation at both the species and population levels, and the connections between different populations and different species, all at once.

The objectives of this report are to assess the importance of accounting for intraspecific genetic correlations in ecological studies. To achieve this, we use the phylogenetic mixed model (PMM) that allows multiple levels of genetic structure to be considered simultaneously (Lynch, 1991; Housworth et al., 2004; Hadfield and Nakagawa, 2010). Other approaches can be used to account for intraspecific correlations (reviewed in Stone et al., 2011; Garamszegi, 2014), but as we show below none currently provides as much flexibility as the PMM. We assess the importance of accounting for intraspecific genetic correlation structure using simulated data and provide a climate change-related empirical example where we investigate the importance of temperature and photoperiod on the timing of budburst of ten tree and shrub species.

## Methods

### The phylogenetic mixed model

The phylogenetic mixed model has been described in detail elsewhere (Housworth et al., 2004; Hadfield and Nakagawa, 2010; Villemereuil and Nakagawa, 2014), thus our description here is brief and focuses on the inclusion of phylogenetic and intraspecific correlations structures as random effects in the model and on the inclusion of fixed effects. In the following, we assume that phylogenetic and intraspecific correlations have been estimated independently, which allows the two structures to be included as separate effects and to quantify their relative importance. Here lowercase italic letters represent numbers, lowercase boldface letters vectors and uppercase boldface letters matrices. The phylogenetic mixed model (PMM) has the form:

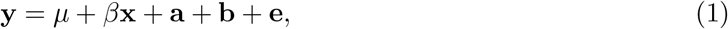

where **y** is the response variable, *µ* is the intercept, **x** is an explanatory variable, *β* the regression coefficient, **a** represents the effects due to the phylogenetic structure, **b** the effects due to the intraspecific structure, and **e** the residuals. **x** is a fixed effect (there could be more than one), whereas **a** and **b** are random effects. The random effects and residuals are assumed to follow normal distributions:

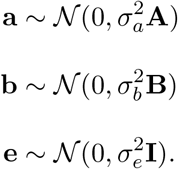

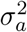 is the phylogenetic variance, 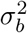 is the intraspecific variance, and 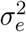 is the residual variance. The matrices **A** and **B** represent the phylogenetic and the intraspecific correlation structures, respectively. The identity matrix **I** indicates that the residuals are independent and identically distributed. Accordingly, the (co)variance structure (**V**) of the model is 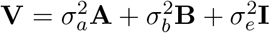.

The total variance 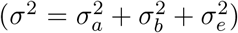 can be partitioned in heritable and non heritable portions. The heritable portion in the present framework consists of the phylogenetic and intraspecific correlation structures. Hence, the heritable proportion of the total variance is 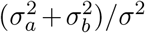. This is akin to the heritability (*h*^2^) parameter in quantitative genetics or the the *λ* parameter in comparative methods (see Housworth et al., 2004, for a discussion). Note that this heritable fraction does not exclusively characterize genetic changes as it can also include non-genetic contributions that can be described by the genetic correlation structures. The remaining variance 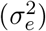, considered non-genetic, includes phenotypic plasticity, measurement error or other effects not defined by the genetic correlation structures.

### Phylogenetic generalized least squares

A brief mention of PGLS seems important as it is a popular comparative method. A PGLS model that would include phylogenetic and intraspecific correlation structures could be denoted as:

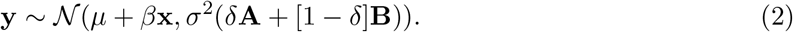

In other words, the residuals of the regression are normally distributed according to a correlation structure that is a combination of phylogenetic and intraspecific effects, with respective weights determined by the parameter *δ*. The main difference with the PMM is the absence of a residual term: in PGLS residuals are completely structured by the genetic correlation matrices provided. This assumption can be relaxed by rescaling the phylogenetic tree to give more or less weight to the terminal branches of the tree (Revell, 2010). We do not consider this PGLS model further here, but it is compared to the PMM using simulations in Appendix S1.

### PMM simulations

Simulations were used to examine the performance of the PMM under a suite of conditions where accounting for intraspecific correlations may be important. We simulated data under the PMM model (equation 1) assuming *µ* = 0 with various relative contributions of the phylogenetic and intraspecific variances, and tested how this affected the estimation of the fixed and random effects. The data simulations followed closely those of Revell (2010) and are described in Appendix S1. One difference is the intraspecific correlation structure that corresponded to the mean variance covariance matrix obtained from 20 independent gene genealogies simulated within a population tree using the Coalescent. The simulations were performed for different regression slopes *β* ∈ {0, 0.1, 0.25} and different ratios of intraspecific and interspecific structure 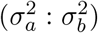 while keeping their sum to 2. In all cases, 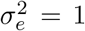 and 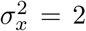. Five hundred simulations were performed for each parameter combination. We simulated data with 98 or 100 individuals but different ratios of species vs. individuals per species, specifically 7 : 14, 10 : 10 and 14 : 7. Larger datasets of 20 species and 20 individuals per species were also simulated.

### Model fitting and performance

The PMM model was fitted using the `MCMCglmm` package in R (Hadfield, 2010). The phylogenetic structure was included in the model by giving the phylogeny to the pedigree argument. The intraspecific structure was incorporated using the genetic intraspecific correlation matrix with a singular value decomposition approach as described in Stone et al. (2011). We used the default priors for the fixed effects and diffuse inverse-Wishart priors for the random effects with *V* = 1 and *ν* = 0.002. We fitted the following models, named according to their respective random effects: 1) *null* (with no genetic structure, **y** = *µ* + *β***x** + **e**); 2) *inter* (phylogenetic structure only, **y** = *µ*+*β***x**+**a**+**e**); 3) *intra* (intraspecific genetic structure only, **y** = *µ*+*β***x**+**b**+**e**); 4) *inter*+*intra* (phylogenetic and intraspecific genetic structures, **y** = *µ* + *β***x** + **a** + **b** + **e**). The MCMC chains were run for 2100 generations, removing the first 100 as burnin and sampling the chain every 10 generations. These settings provided good convergence for all models and all simulation parameters.

Models were compared for accuracy and precision with regards to the estimation of the fixed effects and for the heritable portion of the variance. Accuracy measures how close the estimated slope is to the true value and precision represents the standard deviation of the estimated slope in each MCMC run. We also report the number of simulations that gave a posterior probability > 0.95 for the slope to be greater than 0; this estimates the power of the model when *β* > 0 and the type I error when *β* = 0.

### Budburst experiment

We investigated the usefulness of using the PMM on an ecological study design that included both inter- and intraspecific variation. We analyzed a subset of a larger experiment for which we additionally sampled material for genetic analysis. The experiment’s objective was to determine the impact of temperature increases and longer photoperiods on the budburst timing for several tree and shrub species (full experiment described in Flynn and Wolkovich, 2018). Clippings from 10 species (see Appendix S1) were collected from five individuals at two sites: Harvard Forest (MA, USA; 42.5 °N, 72.2 °W) and the *Station de biologie des Laurentides* (St. Hippolyte, QC, Canada; 45.9 °N, 74.0 °W). Clippings were collected in January 2015 and kept cold until the start of the experiment. They were then subjected to different temperatures (15 °C or 20 °C) and photoperiods (8 or 12 hours of light per day) in growth chambers at the Arnold Arboretum. The number of days to budburst was recorded for each clipping.

To model the interspecific structure, we pruned a published phylogenetic tree of 32,223 angiosperm species based on 7 genes (Zanne et al., 2014) and included it in the model as in the simulations. The intraspecific genetic correlation structure was estimated separately for each species from thousands of genome-wide Genotyping-by-Sequencing markers per species (Elshire et al., 2011). Genetic similarities between individuals within species were estimated using genpofad (Joly et al., 2015), converted into species variance co-variance matrices, and incorporated in the model with a block diagonal matrix as described for the simulations. Details on our methods and R code are provided in Appendices S1 and S2.

The data was analyzed in `MCMCglmm` with warming, photoperiod, and their interaction as fixed effects and time to budburst as the response variable. For the random effects, we fitted the four models used in the simulations in terms of variance structure. We used the same priors as for the simulated data, but ran the chains for 100,000 generations after a burnin of 5000 generations, sampling every 20 generations. MCMC run convergence was assessed using the potential scale reduction factors (PRSF; converges to 1 with increasing convergence). We used the deviance information criterion (DIC) to compare models. Data and commented scripts to replicate the analyses are presented in Appendix S2.

## Results

### Simulations

The *null* model without genetic correlation structure always performed worst in terms of precision and accuracy, whereas the *intra* and *inter* + *intra* models performed best (Fig. 1). The *inter* model with only phylogenetic structure did not perform as well as models *intra* and *inter* + *intra*, but its performance improved with increasing relative importance of the phylogenetic structure over the intraspecific structure. All models performed better when the intraspecific structure was less important. To estimate the heritable proportion of the total variance, the *inter* + *intra* model was the most accurate (Fig. 1), although it slightly overestimated the genetic contribution for greater contributions of intraspecific structure. The *inter* model underestimated the genetic structure of the data, but its estimates were greater than the strict interspecific variance included in the simulations. The *intra* model overestimated the total genetic structure of the data even though only the intraspecific structure was modelled.

**Figure 1:**
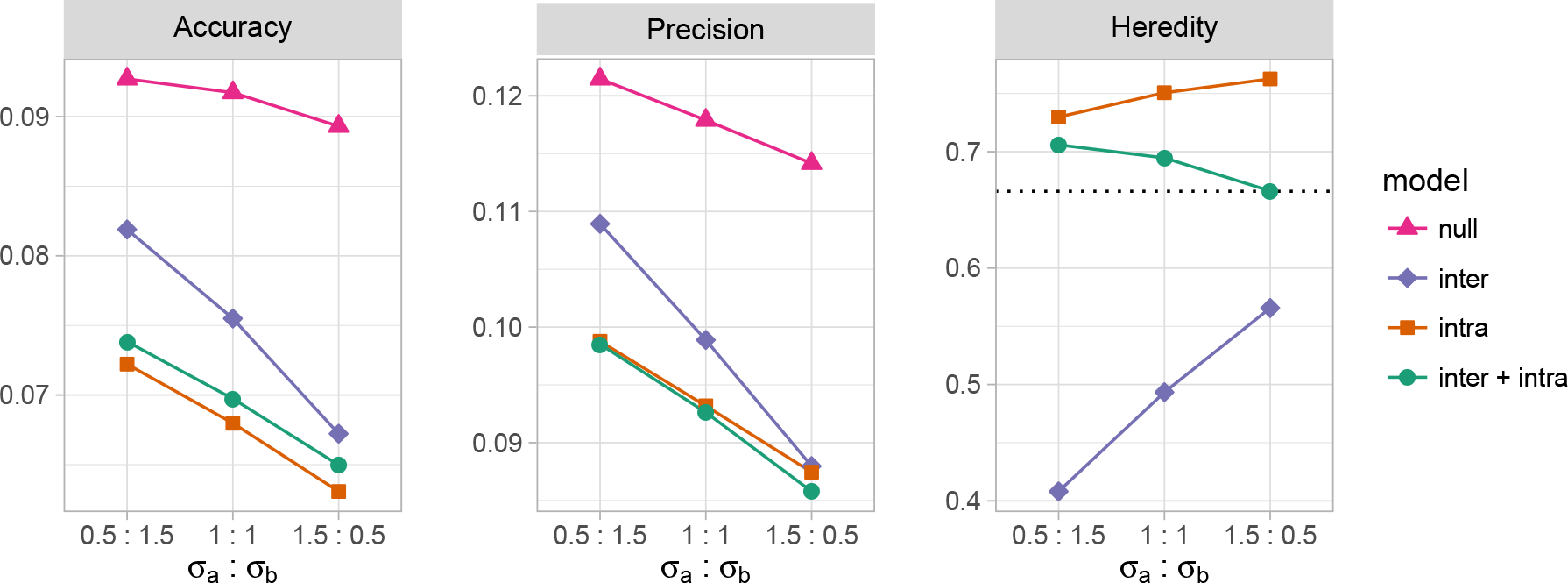
Results of the simulation study for the four variance structure models in terms of slope accuracy and precision, and for estimates of the heritable proportion of the total variance (heredity) with 10 species and 10 individuals per species. Accuracy is the mean absolute distance between the estimated slope 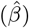 and the true slope (*β*), precision is the mean of the standard deviation of the posterior distribution of 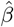 for each simulation, and heredity is the proportion of the total variance explained by the genetic correlation structure (the dashed line indicates the true value). The x-axis indicates the ratio of phylogenetic 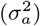 to intraspecific 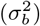 variances used in the simulations. Only the results for *β* = 0.25 are shown as these results were not influenced by the slope.

All models had similar type I error rates (Fig. 2), except for the *intra* model that was slightly higher, especially for increasing importance of the phylogenetic structure. The power of the models was similar for *β* = 0.1, whereas the models *intra* and *inter*+*intra* had the best power for *β* = 0.25.

**Figure 2:**
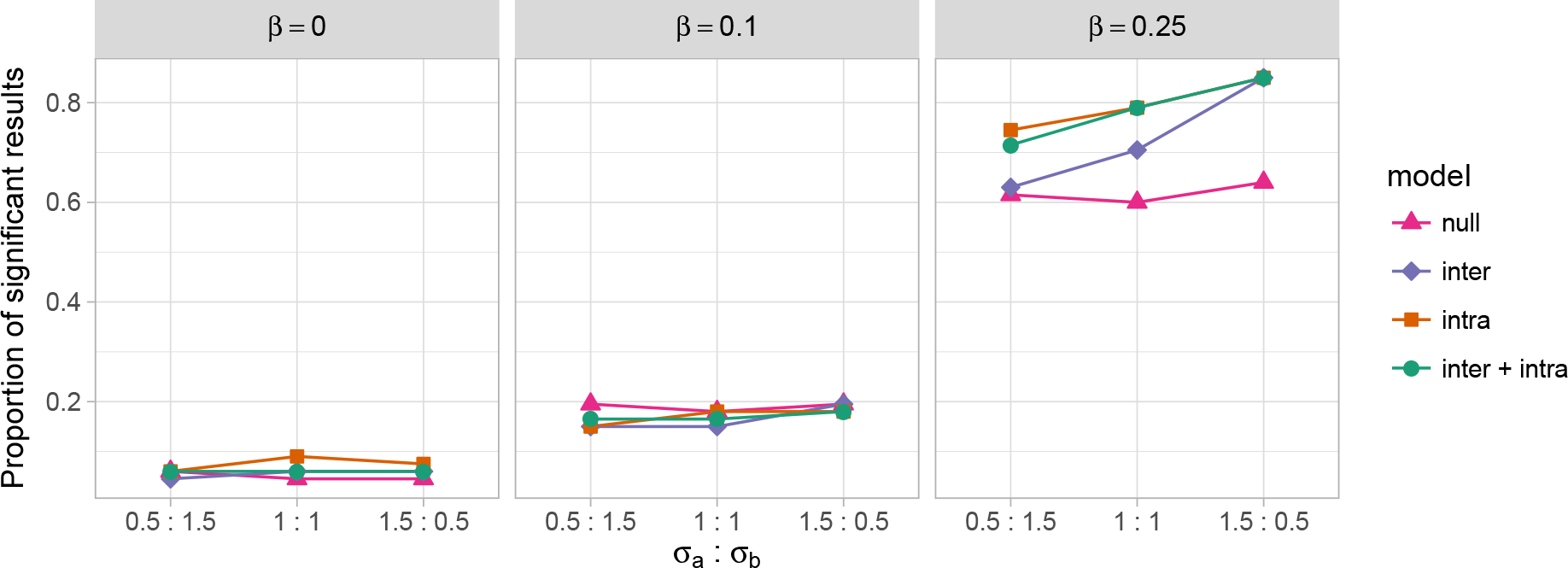
Proportion of the simulations that resulted in a significant regression slope 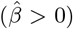 using a threshold of *α* = 0.05. The results for *β* = 0 represent the type I error of the models whereas the results with *β* ∈ {0.1, 0.25} represent the power of the models.

The power of model *inter* with *β* = 0.25 improved with increasing importance of the phylogenetic effect and approached the performance of *intra* and *inter* + *intra* when the phylogenetic effect was three times as important as the intraspecific effect (i.e., when 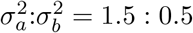).

Varying the amount of population structure had little impact on the results (Appendix S1; Figs. S1, S2). In contrast, increasing the ratio of the number of species to the number of individuals per species resulted in an improved relative performance of the *inter* model compared to models that included the intraspecific structure, but mostly in terms of accuracy (Appendix S1; Figs. S3, S4). Importantly, the advantage of taking into account intraspecific genetic correlations was also present when data were simulated for a single species (Figs. S5, S6). Finally, increasing the samples sizes in the simulations resulted in increased accuracy, precision and power for all methods but did not affect their relative performances (Figs. S7, S8).

### Budburst data

The number of loci obtained per species ranged from 264 in *Prunus* to 2188 in *Vaccinium* and was broadly correlated with the genome size of species (see Appendix S1 for detailed information on the genetic data). The mean locus-based population structure (Φ_*ST*_) between sites was similar across species and ranged from 0.10 to 0.19, suggesting a moderate population structure. Similarly, phylogenetic trees built from the genetic distances showed that individuals from one site were generally more similar to individuals from the same site than to individuals from the other site (Appendix S1).

The MCMC runs showed good convergence (PRSF = 1 for fixed and random effects). The model that best fitted the data was *intra* according to the DIC (2117), followed by *inter* + *intra* (2123), *inter* (2135) and *null* (2426). Incorporating intraspecific structure thus resulted in an important improvement in fit (models *intra* and *inter* + *intra*), while not accounting for genetic correlation (*null*) clearly resulted in a poorer fit.

The wider posterior intervals obtained for the fixed effects with the *null* model illustrate the importance of taking into account the genetic structure present in the data (Fig. 3). This was particularly important for the interaction between warming and photoperiod: the confidence interval included 0 for the *null* model but not the three other models. The three models that accounted for the genetic correlation structure gave similar results, but there was a slight improvement in precision when the intraspecific genetic structure was included. The results suggest that the 5°C warming treatment had the strongest effect on budburst, followed by a longer (four hours) photoperiod (Fig. 3). The interaction between these effects was positive and significant (except for the *null* model), suggesting that they are not additive.

**Figure 3:**
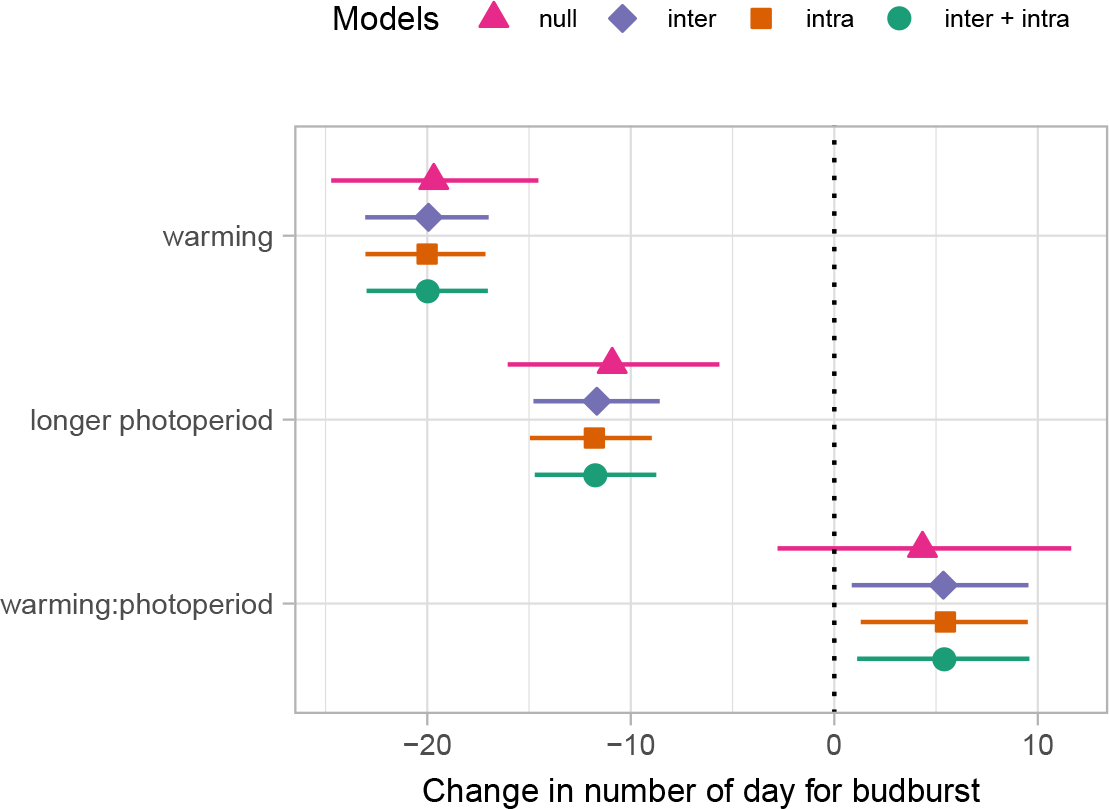
Fixed effects obtained in the phylogenetic mixed model for the four models tested. The symbols represent the means of the posterior distributions and the lines the 95% posterior intervals. The x-axis indicate the change in number of days for budburst, with negative number indicating that buds are opening earlier.

Regarding the partitioning of the variance, the genetic correlation structures explained about two-thirds of the total variance for all models (Table 1). The *inter* + *intra* model that partition the genetic variance into phylogenetic and intraspecific further suggests that the intraspecific variance is slightly greater than the interspecific variance, but the confidence interval was huge suggesting these variances components are difficult to estimate with precision in this dataset (Table 1).

**Table 1:** Mean proportion of the total variance explained by the random effects of the models fitted to explain change in days for budburst, with their 95% posterior intervals (in brackets). The heredity (*h*^2^) and the proportion of the total genetic structure due to the intraspecific correlation 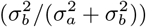 are given for models where they can be estimated.

| Models | $\sigma_a^2$ | $\sigma_b^2$ | $\sigma_e^2$ | $h^2$ | $\sigma_b^2/(\sigma_a^2 + \sigma_b^2)$ |
| --- | --- | --- | --- | --- | --- |
| *null* | – | – | 1 [1,1] | – | – |
| *inter* | 0.65 [0.44, 0.85] | – | 0.35 [0.15,0.56] | 0.65 [0.44, 0.85] | – |
| *intra* | – | 0.68 [0.30, 0.94] | 0.32 [0.06,0.70] | 0.68 [0.30, 0.94] | – |
| *inter + intra* | 0.21 [$4 \times 10^{-6}$, 0.79] | 0.47 [$1 \times 10^{-5}$, 0.93] | 0.31 [0.05,0.66] | 0.69 [0.34,0.95] | 0.69 [$2 \times 10^{-5}$,0.99] |

## Discussion

### Accounting for the intraspecific genetic correlation structure

An increasing number of studies mention the potential importance of accounting for intraspecific genetic structure in multi-species studies. Our results from simulations and empirical data showed multiple advantages of this approach. Perhaps most importantly, incorporating intraspecific correlation structure in statistical models led to a gain in accuracy and precision of the fixed effects, which are generally the parameters of interest in a study. Both the simulations and the empirical studies highlighted this result. The simulations further showed that this advantage persisted under various conditions including when there were no phylogenetic effects (single-species analyses).

The models incorporating intraspecific genetic correlations performed well because they partition the total variance in the data—that would otherwise be classified mostly to the error term—to the genetic correlation structure(s). This is shown by the higher values of the variance due to heritable effects in models that included intraspecific genetic correlations (Fig. 1; Table 1).

In addition to providing more accurate fixed effects, the finer partitioning of the total variance gives a better understanding of the study system by quantifying the proportion of the variance that is due to the intraspecific genetic structure. Notably, the improved performance observed when incorporating the intraspecific structure is not due to the specific modelling framework used in this study, namely the phylogenetic mixed model. Indeed, the simulations we performed under a PGLS model that incorporates intraspecific structure also showed a marked improvement in performance compared to standard ordinary least squares (Appendix S1; Figs. S7, S8).

One surprising result was the very good performance of the model that included only the intraspecific correlation structure (*intra*), which performed nearly as well as the model that accounted for both phylogenetic and the intraspecific genetic structures (*inter* + *intra*) in terms of accuracy and precision in simulations. This result may lead some researchers to consider including only the intraspecific structure, but we advise against it. First, the relative performance of the *intra* model decreased relative to the *inter* + *intra* model when the importance of the intraspecific correlation structure decreased relative to the phylogenetic variance (Fig. 1). Second, the *inter* + *intra* model provided more precise estimates of the proportion of the total variance that is genetically structured in our simulations (Fig. 1) and more precise estimates of fixed effects across a wide range of parameters. And last, as our fundamental biological understanding of ecological questions often stresses the multilevel nature of individuals within species, we argue it is important to include both structures in analyses.

### The Phylogenetic Mixed Model

The phylogenetic mixed model (PMM) offers more flexibility than other comparative methods (Hadfield and Nakagawa, 2010). One advantage is that it uses a terminology familiar to most ecologists. The phylogenetic and the intraspecific genetic correlation structures are considered “random effects” in the model, similar to how blocks are often treated in a randomized block design. That is, the model assumes that they add variance to the species response in a structured way that can be estimated and removed from the residual error, resulting in improved performance of the model. Further, because the residual variance is estimated by the model (in contrast to other methods, see Hadfield and Nakagawa, 2010), model performance is not affected if the intraspecific correlation structure has little effect on the data; in such cases the estimated variance due to intraspecific structure will simply be small.

Another advantage of the PMM is that it allows modeling several random effects simultaneously (Garamszegi, 2014). In our analyses, the total variance of the model included a phylogenetic fraction, an intraspecific fraction, and a residual fraction. But it would be straight-forward to also add a random effect that could account for measurement error, given the appropriate study design. In contrast, the PGLS approach we introduced only considered a phylogenetic and an intraspecific variance with no residual error, which likely explains the increased Type I error compared to the PMM as residual errors were included in the simulations (Appendix S1).

Finally, the PMM is particularly well suited for experimental studies that include several fixed effects as in the present study. In this regard, this study differs from many comparative studies in that the samples studied were subject to experimental treatments. In such experiments, each species has several values for the response variable; at least one per fixed effect. Although such datasets can be analysed using most comparative methods, which often necessitate duplicating the terminal branches of the phylogeny to have—for each species—one tree tip that matches each observation for the response variable, the analysis is much more intuitive with PMM. For instance, the phylogenetic correlation structure can easily be included in the model by associating the species on the phylogeny to the factor representing the species in the dataset (see Appendix S2).

### Modelling guidelines

The importance of accounting for intraspecific genomic relatedness will depend on the importance of the intraspecific genetic structure. Our results showed that the advantages gained from this approach are more important with greater population structure (obtained with smaller effective population sizes, restricted gene flow and longer divergence times) and when the intraspecific variance has a greater relative importance compared to the phylogenetic variance (provided that the phylogenetic structure is corrected for).

The gain from modelling intraspecific correlation structure also depends on the genetic bases of the traits studied and the relevance of the explanatory variable(s) used. In our example, budburst is known to have strong responses to the environmental cues used in the experiment (warming and photoperiod), suggesting these are important explanatory variables. Yet, budburst has also recently been found to be particularly plastic across populations (Aitken and Bemmels, 2016). It is thus possible that the study of other traits with a stronger genetic basis (e.g., timing of budset) could have resulted in larger improvements when accounting for the intraspecific correlation structure.

On a practical aspect, we used the terminology inter- and intraspecific in this study, but the delimitation between the two genetic correlations structures does not have to be at the species level. The decision should be taken depending on the nature of the study. In some cases, it might be logical to have a genetic structure above and below subspecies, and in others such as in recent species complexes it might be interesting to characterize the genetic correlations between closely related species using genome wide markers to capture the complex mosaic structure of genomes.

Comparative methods are being increasingly used to correct for the phylogenetic non-independence of species in statistical tests, in part because of the ease with which one can obtain a well resolved phylogeny. Our results show that important gains can also be obtained by accounting for the intraspecific genetic structure.

### Data and supplementary information

Additional tables and figures are available in Appendix S1 and commented R scripts in Appendix S2. Raw Genotyping-by-sequencing reads have been deposited in the National Center for Biotechnology Information (NCBI) Sequence Read Archive (SRA) under accession number SRP126957. Further scripts, budburst data and processed sequence data will be deposited in a public archive.

### Author’s contribution

SJ and EW conceived the work, DFBF and EW collected the data, SJ analysed the data, SJ and EW wrote the manuscript. All authors gave final approval for publication.

## Acknowledgements

The authors would like to thank T. Savas, J. Samaha, and H. Eyster for help with field collections of tissue for genetic analyses, and constructive comments by Jonathan Davies, Joe Felsenstein, and an anonymous reviewer.

