## Appendix S1 for "On the importance of accounting for intraspecific genomic relatedness in multi-species studies"

#### On the importance of accounting for intraspecific genetic correlations in multi-species studies

Simon Joly, Dan F. B. Flynn, and Elizabeth Wolkovich

#### Description of PMM Simulations

In this section, we provide more details about the simulations performed in the study. Given the Phylogenetic Mixed Model (PMM):

$$\mathbf{y} = \mu + \beta\mathbf{x} + \mathbf{a} + \mathbf{b} + \mathbf{e}, \tag{S1}$$

we simulated data assuming $\mu = 0$, with various relative contributions of the phylogenetic and intraspecific variances and tested how this affected the estimation of the fixed and random effects. The data simulations followed closely those of Revell (2010). The fixed effects $\mathbf{x}$, assumed to be independent and identically distributed, were simulated as $\mathbf{x} = \sqrt{\sigma_x^2}\mathbf{u}$, where $\sigma_x^2$ is the variance of the trait and $\mathbf{u}$ is a vector of random values sampled from the normal distribution. The random effect $\mathbf{a}$ was simulated as $\mathbf{a} = \text{chol}(\sigma_a^2\mathbf{A})'\mathbf{v}$, where $\text{chol}(\sigma_a^2\mathbf{A})$ denotes the upper triangular Cholesky decomposite (chol) of $\mathbf{A}$ times the phylogenetic variance, and $\mathbf{v}$ is a vector of random normal deviates. Similarly, $\mathbf{b} = \text{chol}(\sigma_b^2\mathbf{B})'\mathbf{w}$, where $\mathbf{w}$ is a vector of random normal deviate. Finally, the residual errors, which are independent and identically distributed, were simulated as $\mathbf{e} = \sqrt{\sigma_e^2}\mathbf{z}$, where $\sigma_e^2$ is the error variance and $\mathbf{z}$ is a vector of random normal deviates. The response variable was obtained using $\mathbf{y} = \beta\mathbf{x} + \mathbf{p} + \mathbf{c} + \mathbf{e}$. The simulations were performed for different regression slopes $\beta \in \{0, 0.1, 0.25\}$ and different ratios of intraspecific and interspecific structure ($\sigma_a^2 : \sigma_b^2$) while keeping their sum to 2 (i.e., 0.5 : 1.5, 1 : 1, 1.5 : 0.5). In all cases, $\sigma_e^2 = 1$ and $\sigma_x^2 = 2$.

##### The genetic correlation structures

The phylogenetic correlation structure was obtained by simulating a species tree with $n_{sp}$ species with a pure birth model and extracting the correlation structure from the tree. The intraspecific correlation structure for $n_{ind}$ individuals for each species corresponded to the mean correlation matrix obtained from 20 independent gene genealogies simulated within a population tree using the Coalescent with the `ms` software (Hudson, 2002). The population tree consisted of three populations without migration with speciations 0.3 and 0.8 coalescent units in the past, resulting in moderate population structure. The global intraspecific correlation structure consisted of a block diagonal matrix with the intraspecific correlation matrix of all species along the diagonal. The simulations were performed in `R` (R core team, 2015) and used functions from the packages `phytool` (Revell, 2012), `ape` (Paradis et al., 2004), `phyclust` (Chen, 2011) and `Matrix` (Bates and Maechler, 2016). Five hundred simulations were performed for each parameter combination. We simulated data with 98 or 100 individuals but different ratios of species vs. individuals per species ($n_{sp} : n_{ind}$), specifically 7 : 14, 10 : 10 and 14 : 7.

#### Additional simulations

In this section, we describe additional simulations that were performed to assess the performance of the models and present their results.

##### Variation in the level of population structure

To assess if the level of intraspecific genetic structure chosen for the simulations affected the results, we also performed simulations with both weaker and stronger intraspecific structures. To do so, gene trees were simulated on population trees with shorter and longer coalescent units that resulted in weaker and stronger differentiation between populations, respectively. For the weak population structure, the populations fused 0.15 and 0.4 coalescent units in the past (compared to 0.3 and 0.8 with the simulations presented in the main text), whereas they merged 0.6 and 1.6 coalescent units in the past for the simulations with strong population structure. These simulations were performed with 10 species and 10 individuals per species, with the same settings as for the main simulations.

The simulation results in terms of accuracy and precision were very similar for the different levels of population structure (Fig. S1), with the $inter + intra$ model perhaps performing slightly better with stronger intraspecific structure relative to the other models. The impact of accounting

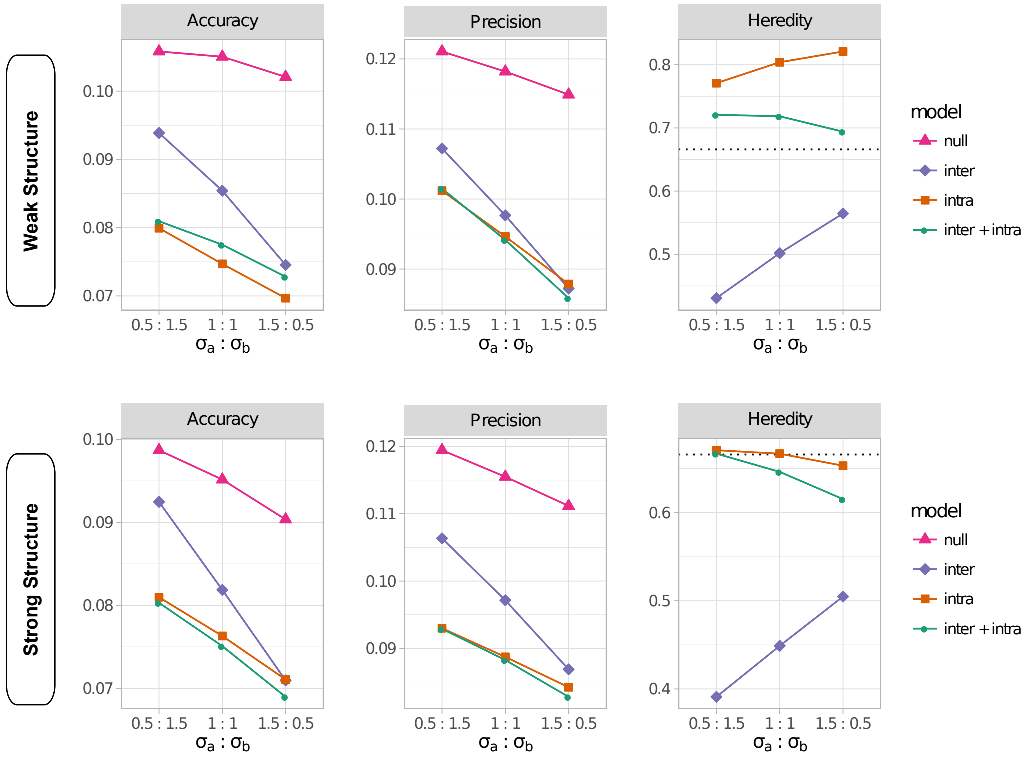

**Figure S1:** Results of the simulation study with the weak and strong population structure settings in terms of slope accuracy and precision, and for estimates of the heritable proportion of the total variance (heredity) with 10 species and 10 individuals per species. Accuracy is the mean absolute distance between the estimated slope ($\hat{\beta}$) and the true slope ($\beta$), precision is the mean of the standard deviation of the posterior distribution of $\hat{\beta}$ for each simulation, and $h^2$ is proportion of the total variance explained by the genetic correlation structure. The x-axis indicates the amount of phylogenetic ($\sigma_a^2$) and intraspecific ($\sigma_b^2$) variances used in the simulations. Only the results for $\beta = 0.25$ are shown as these results were not influenced by the slope.

for the intraspecific population structure was more important for the power of the test, where the stronger population structure resulted in a better relative performance of the models that accounted for the intraspecific genetic structure (Fig. S2).

#### Variation in the number of species and number of individuals per species

We also tested whether the ratio of the number of species to the number of individuals per species affected the simulation results by performing simulations with 7 species and 14 individuals per species, and also 14 species and 7 individuals per species. With these settings, the total size of the dataset was held constant (also compared to the simulation results presented in the main text) and thus only addressed the impact of the ratio of species to individuals per species on the results. Increasing the number of species relative to the individuals per species resulted in a better performance of the *inter* model compared with the models that incorporated intraspecific structure, mostly in terms of accuracy but also for precision and power, although to a lesser level (Figs. S3, S4). But even with the situation with 14 species and 7 individuals per species, the performance of the $inter + intra$ model was still better than the *inter* model for most parameters, except for

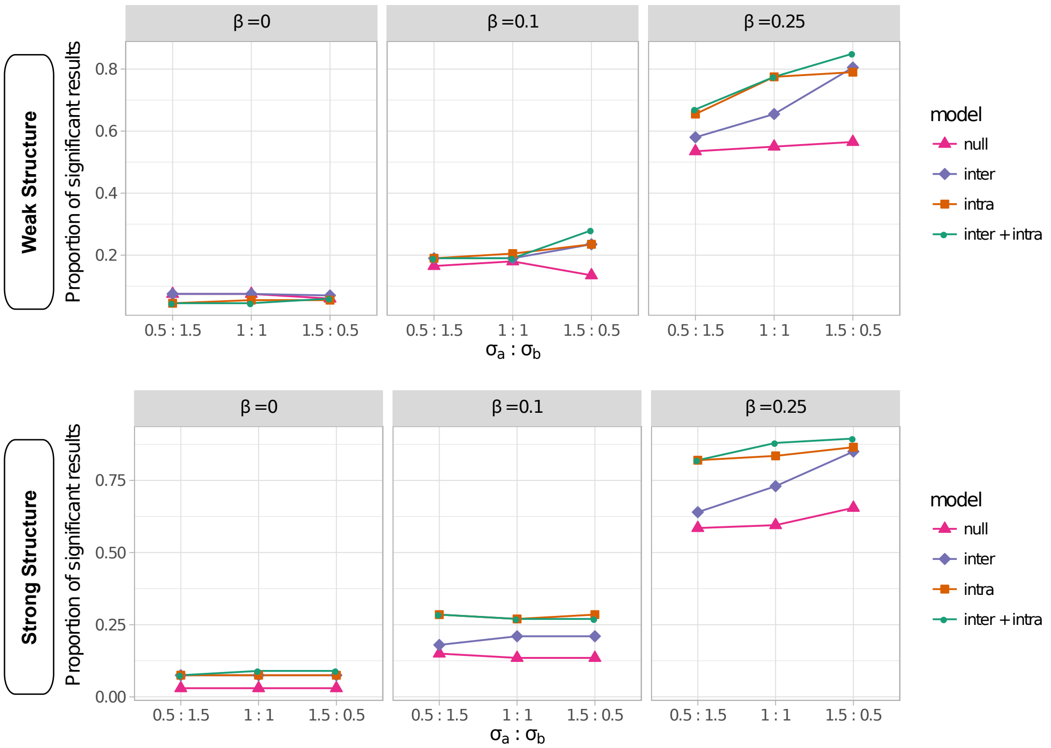

**Figure S2:** Proportion of the simulations that resulted in a significant regression slope ($\hat{\beta} > 0$) using a threshold of $\alpha = 0.05$ when using either weak or strong population structure in the simulations. The results for $\beta = 0$ represent the type I error of the models whereas the results with $\beta \in \{0.1, 0.25\}$ represent the power of the models.

accuracy when the ratio of the phylogenetic to the intraspecific variance was equal or greater than 1 ($\sigma_a \geq \sigma_b$), but only slightly (Fig. S3). In brief, the *inter* + *intra* model performed best in most situations, except where the intraspecific structure was very weak relative to the phylogenetic structure.

#### Simulation with a single species

We performed simulations with a single species to investigate the importance of accounting for intraspecific structure when there was no phylogenetic structure in the data. The simulations proceeded similarly as before, expect that no phylogenetic tree was simulated. One hundred individuals were simulated from three populations with moderate population structure as described for the main simulations. The only difference is that the total variance of the model was made equal to 2 ($\sigma^2 = 2$) in all scenarios tested and we varied the ratio of the intraspecific ($\sigma_b^2$) to the residual error ($\sigma_e^2$). The PMM was fitted using the *null* and the *intra* models.

The results showed that the PMM model accounting for the intraspecific correlation structure resulted in more accurate and more precise slope estimates (Fig. S5). Not surprisingly, the relative performance of the *intra* over the *null* model improved with increasing importance of the intraspecific variance. The Type I error for both models were similar, but the *intra* model resulted in a greater number of significant results compared to the *null* model (Fig. S6). These results

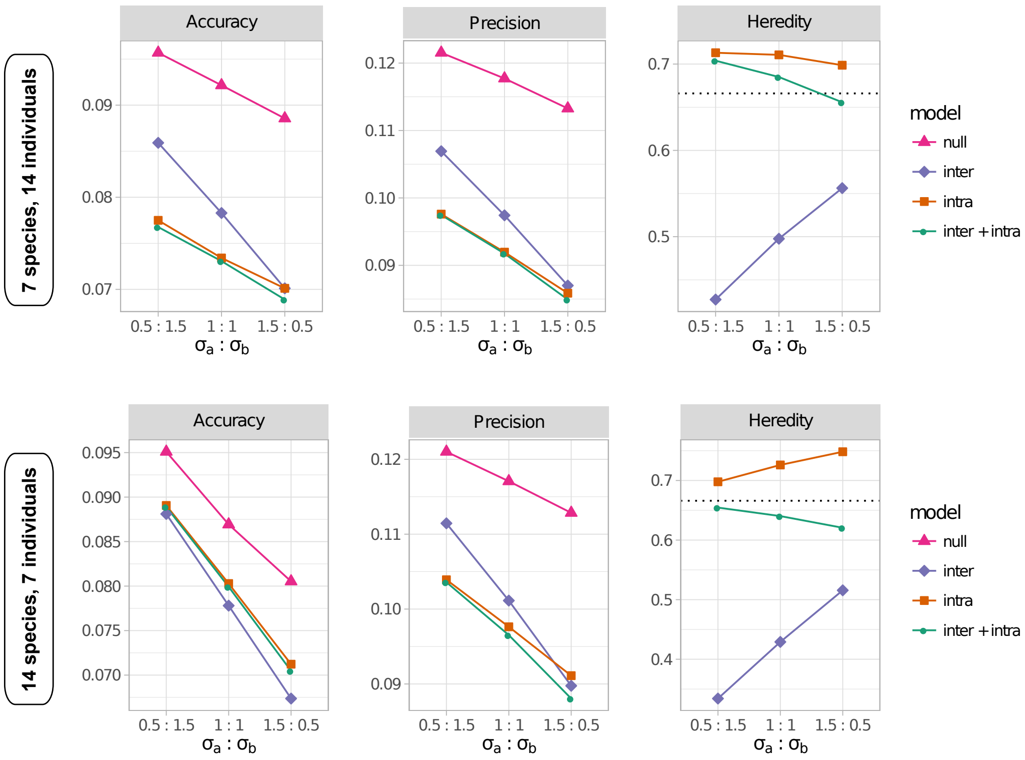

**Figure S3:** Results of the simulation study varying the number of species and individuals per species in terms of slope accuracy and precision, and for estimates of the heritable proportion of the total variance (heredity). Accuracy is the mean absolute distance between the estimated slope ($\hat{\beta}$) and the true slope ($\beta$), precision is the mean of the standard deviation of the posterior distribution of $\hat{\beta}$ for each simulation, and $h^2$ is proportion of the total variance explained by the genetic correlation structure (the dashed line indicates the true value). The x-axis indicates the amount of phylogenetic ($\sigma_a^2$) and intraspecific ($\sigma_b^2$) variances used in the simulations. Only the results for $\beta = 0.25$ are shown as these results were not influenced by the slope.

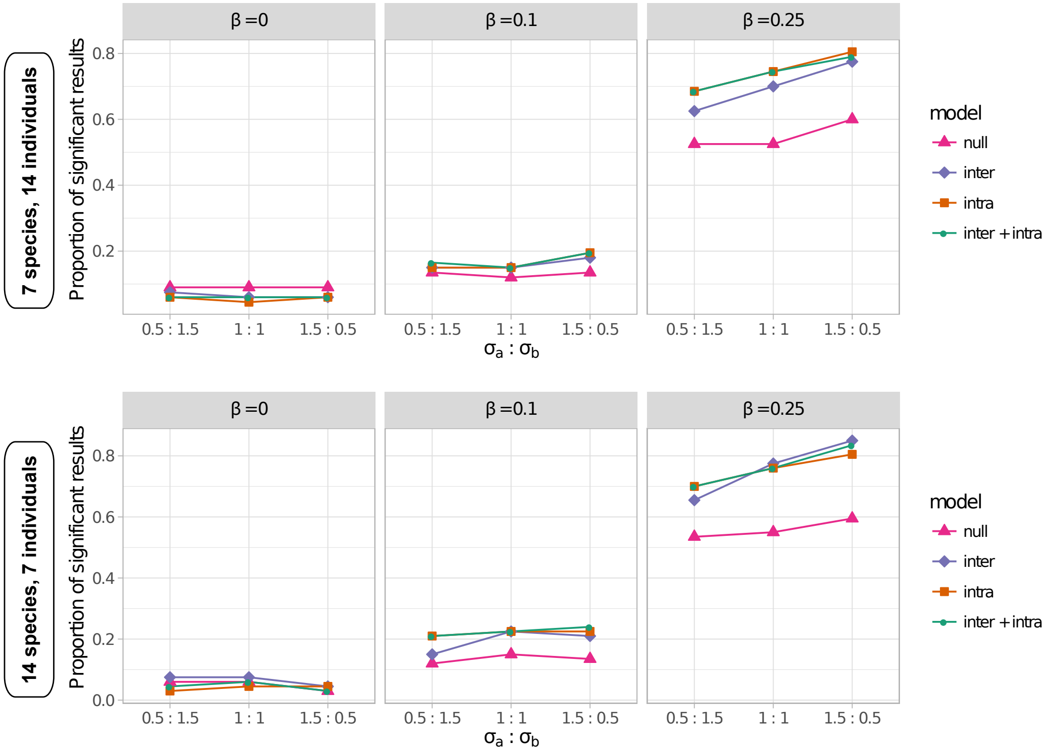

**Figure S4:** Proportion of the simulations that resulted in a significant regression slope ($\hat{\beta} > 0$) using a threshold of $\alpha = 0.05$ when varying the number of species and individuals per species. The results for $\beta = 0$ represent the type I error of the models whereas the results with $\beta \in \{0.1, 0.25\}$ represent the power of the models.

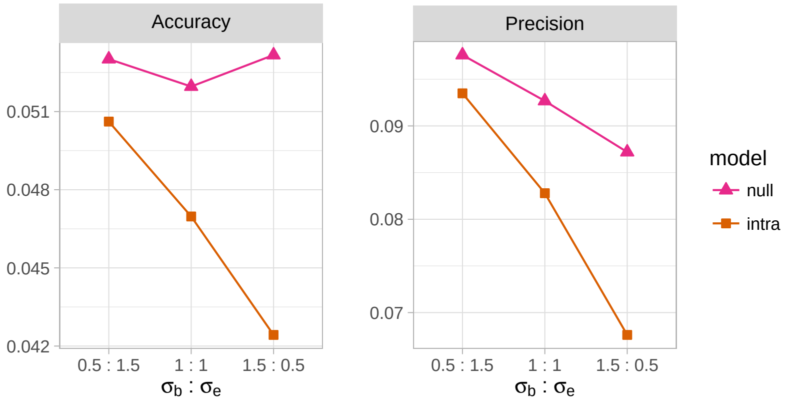

**Figure S5:** Results of the simulation study with a single species in terms of slope accuracy and precision. Accuracy is the mean absolute distance between the estimated slope ($\hat{\beta}$) and the true slope ($\beta$), precision is the mean of the standard deviation of the posterior distribution of $\hat{\beta}$ for each simulation (the dashed line indicates the true value). The x-axis indicates the amount of intraspecific ($\sigma_b^2$) and residual error ($\sigma_e^2$) variances used in the simulations. Only the results for $\beta = 0.25$ are shown as these results were not influenced by the slope.

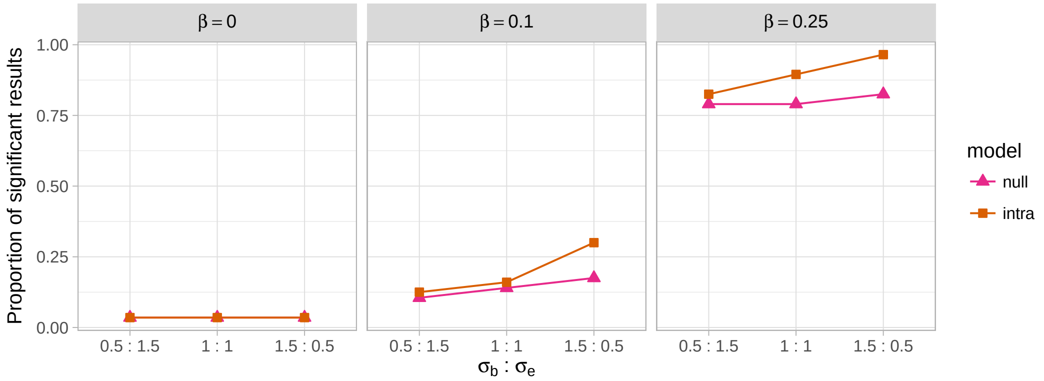

**Figure S6:** Proportion of the simulations that resulted in a significant regression slope ($\hat{\beta} > 0$) using a threshold of $\alpha = 0.05$ in simulations using a single species. The results for $\beta = 0$ represent the type I error of the models whereas the results with $\beta \in \{0.1, 0.25\}$ represent the power of the models.

demonstrate that it is important to account for the intraspecific correlation structure even in the absence of a phylogenetic structure in the data.

#### Increasing the number of species and indiviuals

We performed simulation with twice as much species (20) and individuals per species (20) for a total dataset size of 400 samples. The idea was to investigate how the method behave with greater sample sizes. These simulations showed that all methods were more precise and more accurate with larger sample sizes (Fig. S7). However, the performance of each method relative to the others was not really affected compared with simulations performed with smaller sample sizes. Simulations also showed that all methods showed increased power with greater sample sizes (Fig. S8).

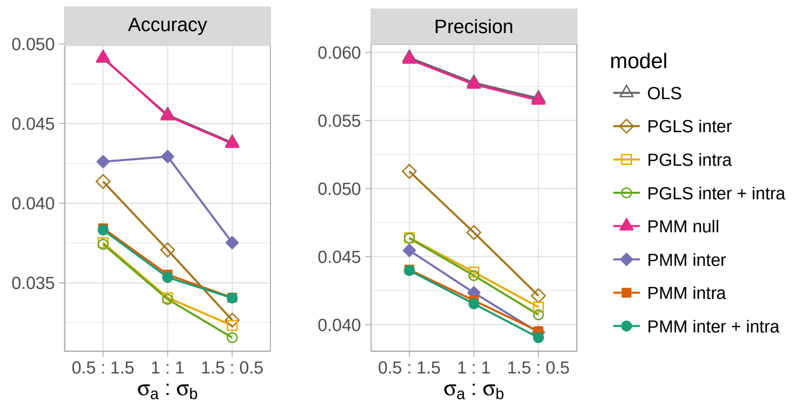

**Figure S7:** Results of the simulation study with 20 species and 20 individuals per species in terms of slope accuracy and precision. Accuracy is the mean absolute distance between the estimated slope ($\hat{\beta}$) and the true slope ($\beta$), precision is the mean of the standard deviation of the posterior distribution of $\hat{\beta}$ for each simulation (the dashed line indicates the true value). The x-axis indicates the amount of intraspecific ($\sigma_b^2$) and residual error ($\sigma_e^2$) variances used in the simulations. Only the results for $\beta = 0.25$ are shown as these results were not influenced by the slope.

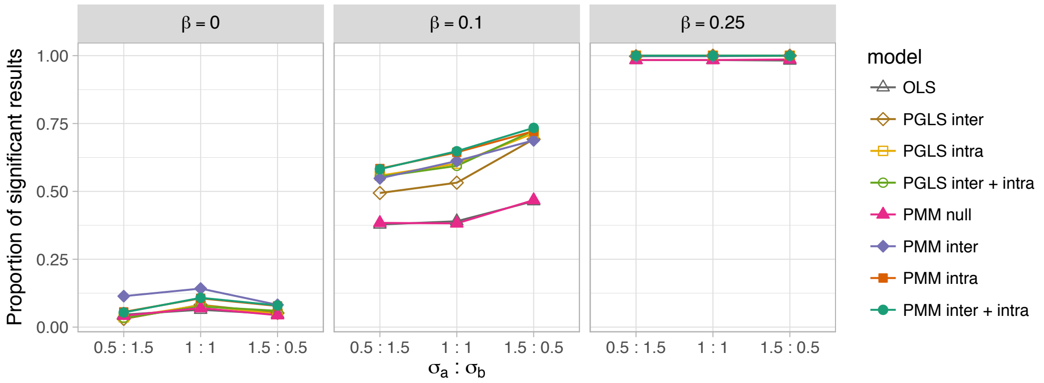

**Figure S8:** Proportion of the simulations that resulted in a significant regression slope ($\hat{\beta} > 0$) using a threshold of $\alpha = 0.05$ in simulations with 20 species and 20 individuals per species. The results for $\beta = 0$ represent the type I error of the models whereas the results with $\beta \in \{0.1, 0.25\}$ represent the power of the models.

### Implementation of a PGLS approach

Phylogenetic generalized least squares (PGLS) is a very popular tool to account for phylogenetic structure in statistical tests. In this section, we describe an approach that corrects simultaneously for both intraspecific and phylogenetic genetic structures in a PGLS framework and compare its performance to the phylogenetic mixed model (PMM) using simulations.

#### Model description

As with the PMM approach, we assume that the phylogenetic and the intraspecific correlation structures have been obtained independently. We propose to combine these two correlation matrices into a global correlation matrix $\mathbf{C}$ that could be included in a generalized least squares approach. Generalized least squares assume that the residuals of a regression $\mathbf{e}$ are distributed according to:

$$\mathbf{e} \sim N(0, \sigma^2 \mathbf{C}).$$

As you can see, all the residual variation is expected to have a mean of 0 and a variance correlated relative to the global genetic correlation matrix. This global genetic correlation matrix consists of a weighted mean correlation structure between the phylogenetic and the intraspecific structures:

$$\mathbf{C} = \delta \mathbf{A} + [1 - \delta]\mathbf{B}.$$

The parameter $\delta$ is the weight and measures the relative importance of the phylogenetic versus intraspecific correlation structure. A greater $\delta$ implies that the phylogenetic correlation structure is more important in describing the variation in the residuals of the model.

More specifically, let $k$ be the number of species, let $i$ and $j$ be species indices, and let $n_i$ be the number of individuals in species $i$. The global correlation matrix ($\mathbf{C}$) could be defined such as

$$\mathbf{C} = \delta \begin{pmatrix} (1)_{n_1 n_1} & \mathbf{P}_{n_1 n_2} & \cdots & \mathbf{P}_{n_1 n_k} \\ \mathbf{P}_{n_2 n_1} & (1)_{n_2 n_2} & \cdots & \mathbf{P}_{n_2 n_k} \\ \vdots & \vdots & \ddots & \vdots \\ \mathbf{P}_{n_k n_1} & \mathbf{P}_{n_k n_2} & \cdots & (1)_{n_k n_k} \end{pmatrix} + (1 - \delta) \begin{pmatrix} \mathbf{W}_{n_1 n_1} & (0)_{n_1 n_2} & \cdots & (0)_{n_1 n_k} \\ (0)_{n_2 n_1} & \mathbf{W}_{n_2 n_2} & \cdots & (0)_{n_2 n_k} \\ \vdots & \vdots & \ddots & \vdots \\ (0)_{n_k n_1} & (0)_{n_k n_2} & \cdots & \mathbf{W}_{n_k n_k} \end{pmatrix}, \tag{S2}$$

where $(1)_{n_i n_i}$ is a square constant matrix of value 1 of size $n_i \times n_i$, $(0)_{n_i n_j}$ is a null matrix of size $n_i \times n_j$, $\mathbf{P}_{n_i n_j}$ is a constant matrix of dimensions equal to $n_i \times n_j$ whose elements are the interspecific correlation $p_{ij}$ between species $i$ and $j$, and $\mathbf{W}_{n_i n_i}$ is a symmetric square correlation matrix of dimension $n_i \times n_i$ containing the intra-specific correlation structure for the $n$ individuals of species $i$. The diagonal of $\mathbf{W}_{n_i n_i}$ is always equal to 1 such that the correlation of an individual with itself is always 1.

In equation (S2), the block matrix at the left of the plus sign determines the inter-specific correlation structure, whereas the right block matrix determines the intra-specific correlation structure. In the inter-specific correlation structure, the matrices of 1s on the diagonal indicate that the individuals within species have a perfect correlation. This make sense as this was not estimated in the inter-specific matrix. In contrast, the non-diagonal matrices of the intra-specific correlation matrix are null matrices, indicating that the inter-specific correlation are 0 for the intra-specific matrix. In practice, each intra-specific correlation matrices $\mathbf{W}_{n_i n_i}$ can be estimated separately for each species, although they should be of the same scale, and can be combined into a diagonal bloc matrix to obtain an intra-specific correlation matrix.

$\delta$ can vary from 0 to 1 and greater values indicate greater importance of the inter-specific correlations in the global genetic correlation matrix ($\mathbf{C}$). A value of 0 would eliminate the left side of the sum and thus remove all inter-specific correlations to leave only intra-specific correlations, whereas a value of 1 would have the opposite effect and delete the intra-specific correlations from the equation. The value of $\delta$ can be given if known. However, in practice, such a scaling parameter might not be know, especially if the phylogenetic and the intraspecific correlation structure are estimated from markers with different and unknown mutation rates. In such cases, $\delta$ could be estimated by generalized least squares along with the other parameters of the model. This optimal $\delta$ should then provide an estimation of the relative importance of the inter-specific correlation matrix over the intra-specific genetic correlation structure for a given dataset.

#### Simulations

We used the same simulation settings as those used in the main manuscript to compare this PGLS approach with the PMM approach. We compared the PGLS approach described above with ordinary least squares (OLS) that do not correct for genetic correlations in the residuals of the model. Moreover, to facilitate comparisons with the results from the Phylogenetic Mixed Model (PMM), we also fitted PGLS models with only phylogenetic structure or only instraspecific structure by fitting $\delta$ to 1 or 0, respectively. The models were fitted using the `gls` function of the `nlme` package in `R` (Pinheiro et al., 2016). We developed a correlation structure called `corIntra` for R, based on the correlation structures available in the `ape` package, to implements the correlation structure we described.

#### Results

The *PGLS inter + intra* approach performed much better than the *OLS* approach in terms of accuracy and precision of the slope (Fig. S9), supporting the findings observed for the PMM. It also performed better than the *OLS inter* and the *OLS intra*. When comparing the *PGLS* models to the PMM models, the *PGLS* models did not perform as well as the PMM models in terms of accuracy, but performed better in terms of precision (Fig. S9). Note, however, that the precision of the *PGLS* is not estimated the same way as that of the the PMM models that are estimated using a Bayesian framework.

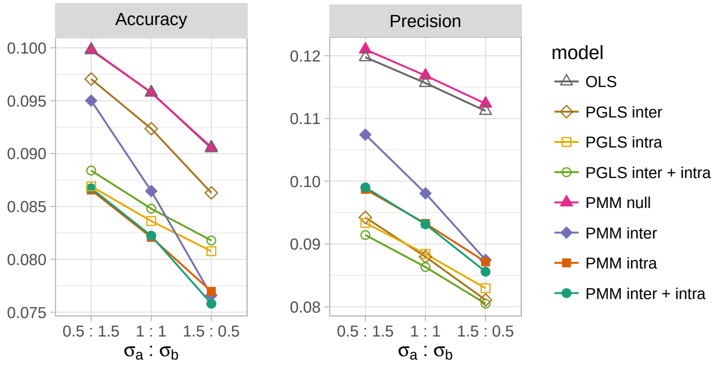

**Figure S9:** Results of the simulation study that included *PGLS* and *OLS* approaches in terms of slope accuracy and precision, and for estimates of the heritable proportion of the total variance (heredity) with 10 species and 10 individuals per species. Accuracy is the mean absolute distance between the estimated slope ($\hat{\beta}$) and the true slope ($\beta$), precision is the mean of the standard deviation of the posterior distribution of $\hat{\beta}$ for each simulation, and $h^2$ is proportion of the total variance explained by the genetic correlation structure. The x-axis indicate the amount of phylogenetic ($\sigma_a^2$) and intraspecific ($\sigma_b^2$) variances used in the simulations. Only the results for $\beta = 0.25$ are shown as these results were not influenced by the slope.

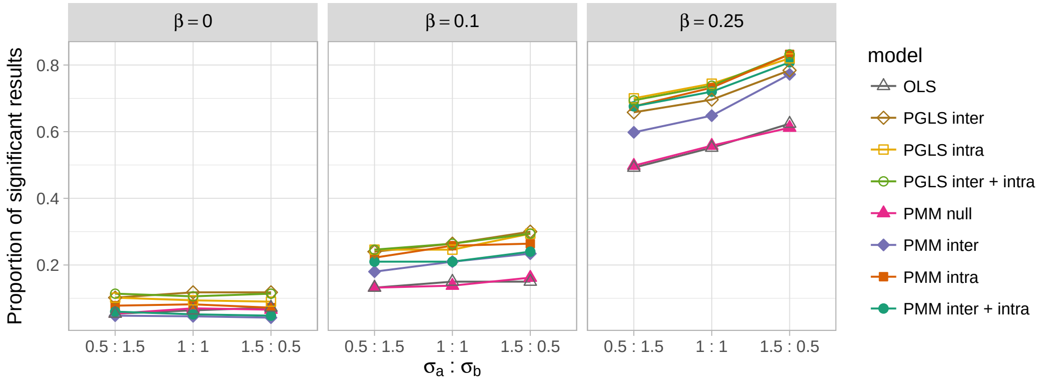

**Figure S10:** Proportion of the simulations that resulted in a significant regression slope ($\hat{\beta} > 0$) with *PGLS*, *OLS* and PMM approaches using a threshold of $\alpha = 0.05$. The results for $\beta = 0$ represent the type I error of the models whereas the results with $\beta \in \{0.1, 0.25\}$ represent the power of the models.

The *PGLS* models had similar power than their equivalent PMM models, or slightly better. However, the *PGLS* models also had an increased type I error (Fig. S10). This could probably be attributed to the fact that PGLS assume that all the residuals correlation is genetically structured, whereas this was not the case in the simulations. Overall, the PMM seem to have a slightly better performance over the PGLS approach to simultaneously model phylogenetic and intraspecific genetic correlation structures. It might be possible to further improve the performance of the PGLS approach by adding an additional parameter that would allow the residuals to be less correlated, similar to the $\lambda$ parameter often used in PGLS (Revell, 2010), but investigating this is beyond the scope of the present study.

### Genetic analyses of the budburst dataset

#### Supplementary methods

##### DNA extraction and library preparation and sequencing

Leaves were immediately placed in presence of silica gel on the field. The DNA was extracted using the plant DNA kit from QIAGEN (Mississauga, Ontario). DNA concentrations were determined on gel and 200 ng of each sample in 10 $\mu$L was sent to the *Institut de biologie intégrative et des systèmes* (IBIS) of University Laval for Genotyping-by-sequencing (GBS) library preparation. The library was prepared following Elshire et al. (2011) using the enzyme combination SbfI (CCTGCA/GG) – MspI (C/CGG). This combination was chosen to reduce the number of fragments and thus maximize the sequencing depth, a conservative strategy for obtaining good SNP calling since the samples involved several species with various genome sizes (Table S1). The libraries (single ends; 100 bp) were sequenced at the Genome Quebec Innovation Centre (Montreal, Canada) on one lane of a HiSeq 2000 Illumina sequencer.

##### Bioinformatics

The reads were filtered for quality using Trimmomatic vers. 0.35 (Bolger et al., 2014) with the following sequential steps: adapter trimming (seed mismatch = 3; clip threshold = 6), removing leading and trailing nucleotides with phred scores < 15, removing the remaining nucleotides of a read after the mean nucleotide phred score within a sliding window of 5 nucleotides drops below 15, and finally remove all reads < 50 bp. Sequence quality was inspected before and after filtering using FastQC vers. 0.11.4 (http://www.bioinformatics.babraham.ac.uk/projects/fastqc/). The software Stacks vers. 1.35 (Catchen et al., 2013) was used to generate loci de novo and perform SNP calling. Amongst most critical parameters, we set -m to 3, -M to 3, and -max_locus_stacks to 4 in ustacks, and -n to 5 in cstacks. These parameters were chosen following Mastretta-Yanes et al. (2015) and considering that the populations compared are geographically isolated. Most other parameters were set to default. The loci assembly and SNP calling was performed independently for the different species.

##### Analysis of GBS data

Only the loci that were present in at least 60% of the individuals in each population were used in the genetic analyses. We estimated locus-based $F_{ST}$ statistics ($\Phi_{ST}$) with Stacks to quantify the overall population structure. However, to allow a finer description of the genetic structure, we estimated the genetic distances between the individuals from the DNA sequences of the full loci obtained from Stacks. We used the genpofad distance from the `pofadinr R` package (Joly, 2016) that has the advantage of incorporating polymorphic SNPs in distances and that provides an accurate estimate of true genetic distances (Joly et al. 2015). These distances were used to build the infraspecific correlation matrix, but also for estimating neighbour-joining tree (Saitou and Nei 1987) separately

**Table S1:** Haploid genome sizes (1C) for the species included in the study, in mega base pairs (Mbp). In cases where the genome size of the species is not available, the genome size of a close relative of the same policy level is given instead.

| Species | Genome size (1C) | Method | Reference |
|---|---|---|---|
| *Acer pensylvanicum* | 675 Mbp[1] | Flow cytometry | Siljak-Yakovlev et al. (2010) |
| *Alnus incana* | 553 Mbp | Flow cytometry | Siljak-Yakovlev et al. (2010) |
| *Fagus grandifolia* | 528 Mbp | Flow cytometry | Bainard et al. (2011) |
| *Lonicera canadensis* | 929 Mbp[2] | Flow cytometry | Zonneveld et al. (2005) |
| *Populus grandidentata* | 484 Mbp[3] | Flow cytometry | Bennett and Leitch (2012) |
| *Prunus pensylvanica* | 262 Mbp[4] | Flow cytometry | Bennett and Leitch (2012) |
| *Quercus rubra* | 831 Mbp | Flow cytometry | Horjales et al. (2003) |
| *Spirea alba* | – | – | – |
| *Vaccinium myrtilloides* | 616 Mbp | Flow cytometry | Costich et al. (1993) |
| *Viburnum lantanoides* | 3907 Mbp[3] | Flow cytometry | Bennett and Leitch (2012) |

Notes: [1]Value for *A. campestre*; [2]Value for*L. nitida*; [3]Genus mean; [4]Mean of diploid species of the genus.

for each species to see whether the individuals within each populations were more similar with each other than to individuals from the other population.

To build the intraspecific correlation matrix that will be used in the Phylogenetic Mixed Model (PMM), we standardized the distance matrix obtained for each species by dividing each distance by the largest distance in the matrix. We then converted the distances in similarities and created a block diagonal matrix to use in the Phylogenetic Mixed Model.

#### Results

The sequencing resulted in a total of 175,511,015 reads of 100 nucleotides. After cleaning, a total of 172,821,478 reads remained (98.48%). The reads that passed the filter were of good quality and no adapters were detectable in the sequences. In general, the number of reads were relatively well distributed among individuals (Fig. S11).

The number of loci obtained per species ranged from 264 in *Prunus* to 2188 in *Vaccinium* (Fig. S12). These numbers broadly correlate with the genome size of the different species, but not strictly so (Pearson $r = 0.59$, $p = 0.092$; Table S1). Note that few loci were recovered for all individuals in each species (Fig. S12). Nevertheless, the sequencing depth per loci was relatively good (Fig. S13), with a mean depth above 27 for all species, which shows that no individual had a particularly poor sequencing depth.

##### Population structure

Locus based $\Phi_{ST}$ was similar across species with a mean $\Phi_{ST}$ that ranged from 0.10 to 0.19 (Fig. S14). The neighbour-joining trees representing the genome wide similarity among individuals support the important population structure (Fig. S15). For five species out of ten, individuals from the two sites formed distinct groups indicating that individuals were always closer to an individuals from its own population than to an individual from the other population (Fig. S15). For the remaining species, only one individual typically grouped with the other population. There are two exceptions. *Populus grandidentata* doesn't show clear genetic structure among populations and the individuals

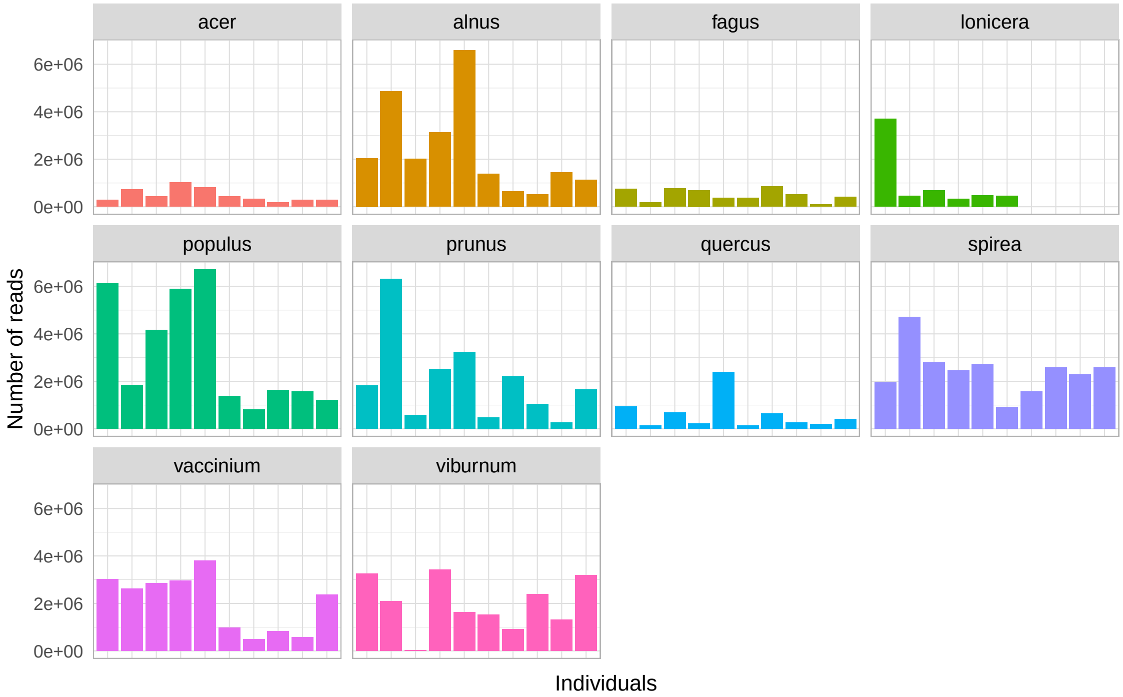

**Figure S11:** Number of reads for each individuals included in the analysis.

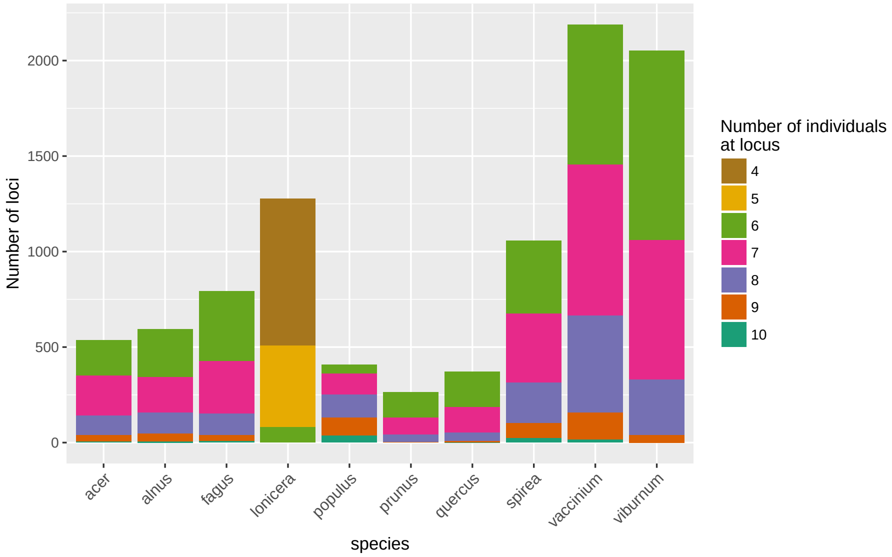

**Figure S12:** Number of loci obtained for each species and number of individuals for which the loci was obtained.

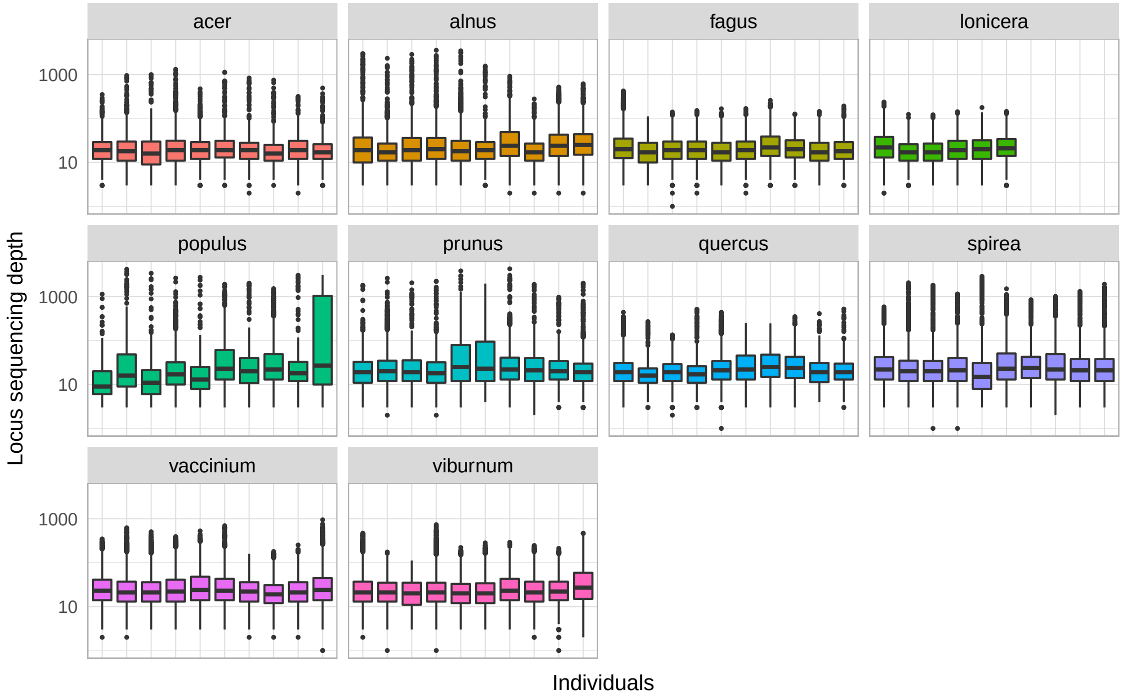

**Figure S13:** Number of sequences per locus obtained for each species.

are much less differentiated than for the other species (Fig. S15). The other exception is *Prunus pensylvanica*, for which two individuals from Massachusetts represent outliers and are more clearly more distant than the remaining individuals (Fig. S15).

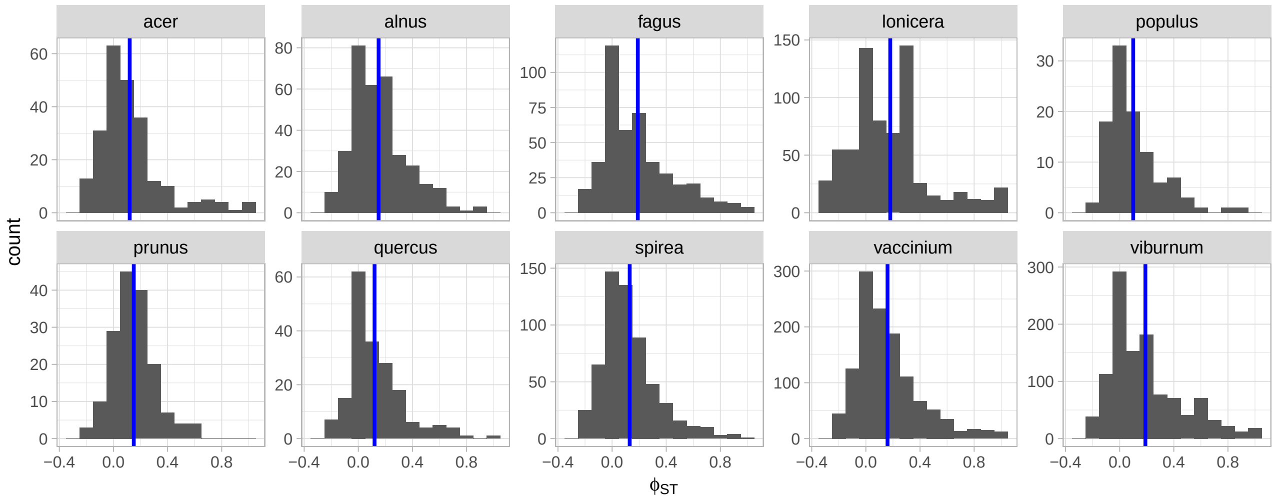

**Figure S14:** Locus-based $\Phi_{ST}$ distributions for all loci between the two sites. The blue lines indicate the mean $\Phi_{ST}$ across the genome.

Costich, D. E., R. Ortiz, T. R. Meagher, L. P. Bruederle, and N. Vorsa. 1993. Determination of ploidy level and nuclear DNA content in blueberry by flow cytometry. Theoretical and Applied Genetics **86**:1001–1006.

Elshire, R. J., J. C. Glaubitz, Q. Sun, J. A. Poland, K. Kawamoto, E. S. Buckler, and S. E. Mitchell. 2011. A robust, simple genotyping-by-sequencing (GBS) approach for high diversity species. PLoS ONE **6**:e19379.

Horjales, M., N. Redondo, A. Blanco, and M. A. Rodríguez. 2003. Cantidades de DNA nuclear en árboles y arbustos. NACC: Nova Acta Científica Compostelana. Bioloxía pages 23–33.

Hudson, R. R. 2002. Generating samples under a Wright-Fisher neutral model of genetic variation. Bioinformatics **18**:337–338.

Joly, S., 2016. pofadinr: Distance methods from SNP data. URL `https://github.com/simjoly/pofadinr`.

Mastretta-Yanes, A., N. Arrigo, N. Alvarez, T. H. Jorgensen, D. Piñero, and B. C. Emerson. 2015. Restriction site-associated DNA sequencing, genotyping error estimation and de novo assembly optimization for population genetic inference. Mol Ecol Resour **15**:28–41.

Paradis, E., J. Claude, and K. Strimmer. 2004. APE: Analyses of Phylogenetics and Evolution in R language. Bioinformatics **20**:289–290.

Pinheiro, J., D. Bates, S. DebRoy, D. Sarkar, and R Core Team, 2016. nlme: linear and nonlinear mixed effects models. URL `http://CRAN.R-project.org/package=nlme`.

R core team, 2015. R: a language and environment for statistical computing. URL `http://www.R-project.org`.

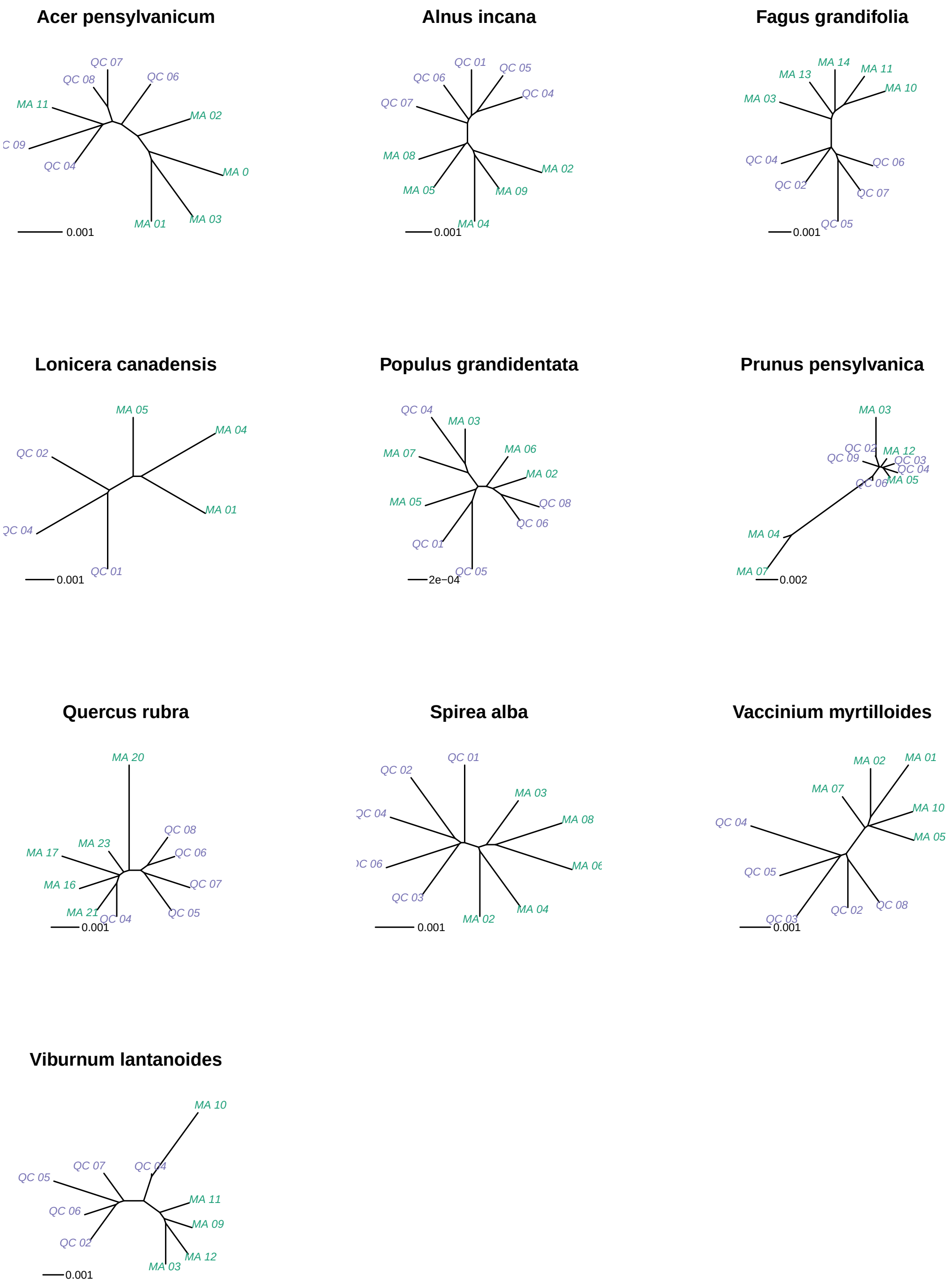

**Figure S15:** Neighbour-joining trees of individuals representing the overall genetic similarities among individuals for all species. The blue individuals are from the Quebec site whereas the green individuals are from the Massachusetts site.

Revell, L. J. 2010. Phylogenetic signal and linear regression on species data. Methods Ecol Evol **1**:319–329.

Revell, L. J. 2012. phytools: An R package for phylogenetic comparative biology (and other things). Methods in Ecology and Evolution **3**:217–223.

Siljak-Yakovlev, S., F. Pustahija, E. M. Šolić, F. Bogunić, E. Muratović, N. Bašić, O. Catrice, and S. C. Brown. 2010. Towards a genome size and chromosome number database of Balkan flora: C-values in 343 taxa with novel values for 242. Advanced Science Letters **3**:190–213.

Zonneveld, B. J. M., I. J. Leitch, and M. D. Bennett. 2005. First nuclear DNA amounts in more than 300 angiosperms. Ann Bot **96**:229–244.
