## Appendix S2 for "On the importance of accounting for intraspecific genomic relatedness in multi-species studies"

### Appendix S2: Analysis of budburst data using MCMCglmm

*Simon Joly and Elizabeth Wolkovich*

*December 2018*

### Preface

This file describe the analysis of the budburst data using Phylogenetic Mixed Models using the MCMCglmm R package.

### Data preparation

#### Load packages

```
# Packages -----
library(ape)
library(MCMCglmm)
library(ggplot2)
library(plyr)
```

#### Prepare the datasets

Let's load the datasets that we will need to fit the models.

```
# Import the data files
load(file="./data/budbust-nonstand.Rdata")
```

There are three objects:

1. `thedata` contains the budburst data. It contains the species information (`sp`), the time to budburst (`lday`), the warming treatment (`warm`), the photoperiod treatment (`photo`), the population from where the individual was collected (`site`), and a code identifying the individuals (`code`).
2. `treephylo` contains the phylogenetic tree
3. `intra.mat` contains the block diagonal matrix of the unstandardized intraspecific genetic similarities.

Let's have a look at the first lines of the data.

```
head(thedata,n=8)
```

```
##                    sp lday warm photo site        code
## ACEPEN01_HF1 ACEPEN   48   15    12   HF ACEPEN_MA_01
## ACEPEN01_HF2 ACEPEN   34   20     8   HF ACEPEN_MA_01
## ACEPEN01_HF3 ACEPEN   26   20    12   HF ACEPEN_MA_01
## ACEPEN01_HF4 ACEPEN   68   15     8   HF ACEPEN_MA_01
## ACEPEN02_HF1 ACEPEN   40   15    12   HF ACEPEN_MA_02
## ACEPEN02_HF2 ACEPEN   68   15     8   HF ACEPEN_MA_02
## ACEPEN02_HF3 ACEPEN   NA   20    12   HF ACEPEN_MA_02
## ACEPEN02_HF4 ACEPEN   40   20     8   HF ACEPEN_MA_02
```

##### Format the data for the MCMCglmm analysis

```
# MCMCglmm data preparation ----

# Single value decomposition of the intraspecific structure
# follow code of Stone et al. 2012
intra.svd <- svd(intra.mat)
intra.svd <- intra.svd$v %*% (t(intra.svd$u) * sqrt(intra.svd$d))
rownames(intra.svd) <- colnames(intra.svd) <- rownames(intra.mat)

# Remove node names from phylogeny
treephylo$node.label <- NULL
# MCMCglmm needs ultrametric trees
is.ultrametric(treephylo)
```

```
## [1] FALSE
```

```
# Make tree ultrametic
treephylo <- chronos(treephylo)
```

```
##
## Setting initial dates...
## Fitting in progress... get a first set of estimates
##          Penalised log-lik = -59.30466
## Optimising rates... dates... -59.30466
## Optimising rates... dates... -59.30403
##
## Done.
```

```
class(treephylo) <- "phylo"
# check again
is.ultrametric(treephylo)
```

```
## [1] TRUE
```

```
# Rename the column with species names "animal". This is required for MCMCglmm
colnames(thedata)[1] <- "animal"

# Prepare intraspecific correlation matrix. Specifically, the individuals have to
# appear the same number of times as in the dataset and in the same order
matches <- match(thedata[,"code"],rownames(intra.svd))
# Reorder intra.svd to fit the data
```

```
intra.svd <- intra.svd[unique(matches),unique(matches)]
# Inflate matrix: duplicate rows and comlums for the number of treatments
# present in the dataset.
intra.svd <- matrix(apply(apply(intra.svd,2,function(c)
  rep(c,times=table(matches))),1,function(c)
    rep(c,times=table(matches))),nrow=sum(table(matches)),ncol=sum(table(matches)))

# Finally, remove rows with missing data
thedata <- na.omit(thedata)
intra.svd <- intra.svd[-attr(thedata,"na.action"),-attr(thedata,"na.action")]
```

### Data analysis

#### Model fitting

Fit different models using MCMCglmm. A specific prior is built for each model.

```
# Model M0 is a simple model with no random factors
priorpr.m0 <- list(R = list(V = 1, nu = 0.002))
M0 <- MCMCglmm(lday ~ warm * photo, data=thedata, scale=TRUE,
               nitt=105000,thin=20,burnin=5000,prior=priorpr.m0)

# Model M1 is a simple phylogenetic model without intraspecific correlation structure
priorpr.m1 <- list(R = list(V = 1, nu = 0.002),
                   G = list(G1 = list(V = 1, nu = 0.002)))
M1 <- MCMCglmm(lday ~ warm * photo,
               random = ~ animal, pedigree = treephylo, data=thedata,
               scale=TRUE,nitt=105000,thin=20,burnin=5000,prior=priorpr.m1)

# Model M2 has only intraspecific structure
M2 <- MCMCglmm(lday ~ warm * photo,
               random =~ idv(intra.svd), data=thedata, scale=TRUE,
               nitt=105000,thin=20,burnin=5000,prior=priorpr.m1)

# Model M3 has both phylogenetic and intraspecific structure
priorpr.m3 <- list(R = list(V = 1, nu = 0.002),
                   G = list(G1 = list(V = 1, nu = 0.002),
                            G2 = list(V = 1, nu = 0.002)))
M3 <- MCMCglmm(lday ~ warm * photo,
               random =~ idv(intra.svd) + animal,
               pedigree = treephylo, data=thedata, scale=TRUE,
               nitt=105000,thin=20,burnin=5000,prior=priorpr.m3)
```

##### Model comparison

Now that we ran the models, we can compare the different models using the Deviance Information Criterion (DIC; lower values are best).

```
# Compare fit of the models (deviance information criterion)
data.frame(models=c("M0","M1","M2","M3"),
           random.effects=c("NA","inter","intra","inter+intra"),
           DIC=c(M0$DIC,M1$DIC,M2$DIC,M3$DIC))
```

```
##   models random.effects      DIC
## 1     M0             NA 2425.909
## 2     M1          inter 2134.531
## 3     M2          intra 2117.913
## 4     M3    inter+intra 2122.693
```

##### Run convergence

Before looking at this model more closely, it is important to look at the convergence of the MCMC run. To evaluate this, it is useful to run an independent analysis. Let's do this for the model 3.

```
# Do another run to check for convergence
M3b <- MCMCglmm(lday ~ warm * photo,
                random =~ idv(intra.svd) + animal,
                pedigree = treephylo, data=thedata, scale=TRUE,
                nitt=105000,thin=20,burnin=5000,prior=priorpr.m3)
```

Once this is done, we can calculate the Potential scale reduction factors for the random effects.

```
# Potential scale reduction factors (PSRF)

# PSRF of fixed effects
gelman.diag(mcmc.list(M3$Sol,M3b$Sol))
```

```
## Potential scale reduction factors:
##
##                Point est. Upper C.I.
## (Intercept)          1.01       1.01
## warm20               1.00       1.00
## photo12              1.00       1.00
## warm20:photo12       1.00       1.00
##
## Multivariate psrf
##
## 1
```

```
# PSRF of random effects
gelman.diag(mcmc.list(M3$VCV,M3b$VCV))
```

```
## Potential scale reduction factors:
##
##            Point est. Upper C.I.
## intra.svd.       1.03       1.11
## animal           1.00       1.01
## units            1.01       1.04
##
## Multivariate psrf
##
## 1.02
```

You can see that these are very close to 1, suggesting good convergence. This is also evident when looking at the plot of the values generation per generation as the mixing is very good.

```
# look at MCMC chain sampling
plot(mcmc.list(M3$VCV,M3b$VCV))
```

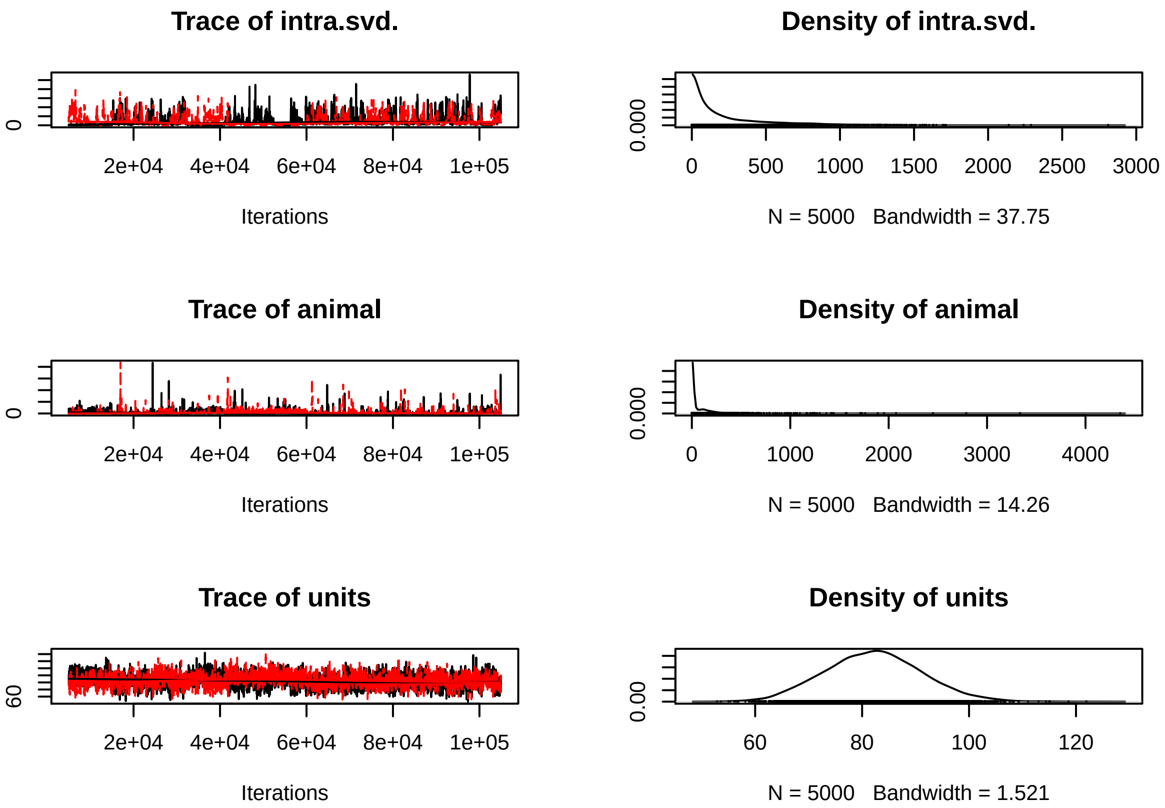

##### Model summary

Now that we are convinced that the analyses have converged, we can look at the results summary (here from a single run).

```
# Summary of best model
summary(M3)
```

```
##
##  Iterations = 5001:104981
##  Thinning interval  = 20
##  Sample size  = 5000
##
##  DIC: 2122.693
##
##  G-structure:  ~idv(intra.svd)
##
##             post.mean  l-95% CI u-95% CI eff.samp
## intra.svd.     237.8 0.0002957    892.7    136.3
##
##                 ~animal
##
##         post.mean  l-95% CI u-95% CI eff.samp
## animal     81.55 0.0001972    325.4    524.3
##
##  R-structure:  ~units
##
##       post.mean l-95% CI u-95% CI eff.samp
```

```
## units     81.97    64.39    99.66    257.8
##
##  Location effects: lday ~ warm * photo
##
##                  post.mean l-95% CI u-95% CI eff.samp  pMCMC
## (Intercept)         57.476   38.765   78.918     3776 <2e-04 ***
## warm20             -19.977  -22.980  -17.024     5000 <2e-04 ***
## photo12            -11.731  -14.716   -8.747     5000 <2e-04 ***
## warm20:photo12       5.418    1.124    9.585     5000 0.0136 *
## ---
## Signif. codes:  0 '***' 0.001 '**' 0.01 '*' 0.05 '.' 0.1 ' ' 1
```

The effective sample sizes (corrected for the autocorrelation along the chain) are all relatively good, both for random and fixed effects.

We can also visualize the fixed effects using a figure.

```
#Fixed effects
results.M3 <- as.data.frame(summary(M3)$solutions)
results.M3$effects <- factor(rownames(results.M3),levels=rownames(results.M3))
colnames(results.M3)[2:3] <- c("lowerCI","upperCI")
fixed.p <- ggplot(results.M3[-1,], aes(y=post.mean, x=effects)) +
  geom_point(color="black") +
  geom_linerange(aes(ymin=lowerCI, ymax=upperCI)) +
  geom_hline(yintercept = 0, lty=3) +
  scale_x_discrete(limits = rev(levels(results.M3$effects)[-1])) +
  ylab("Change in number of day for budburst") + xlab("") +
  coord_flip() + theme_light()
fixed.p
```

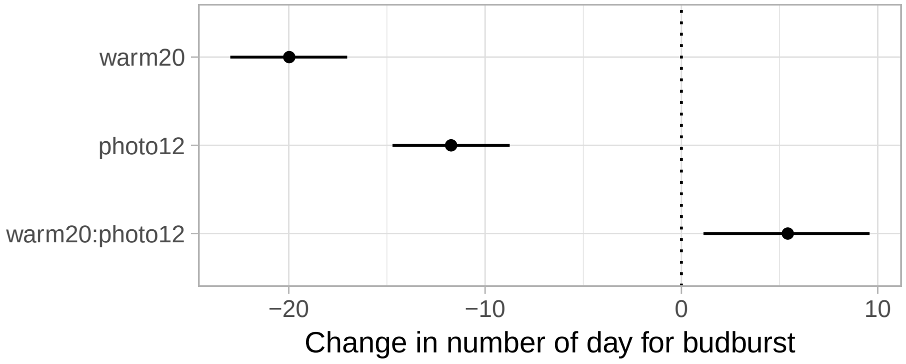

For the **fixed effects**, the model suggests that temperature and the photoperiod are highly significant. The interaction between the temperature and the photoperiod (`warm:photo`) is also significant.

Let's have a look at the raw data:

```
library(ggplot2)
thedata$photo.temp <- interaction(thedata$photo,thedata$warm)
p <- ggplot(thedata,aes(y=lday,x=site))
p + geom_boxplot(aes(fill=photo.temp),outlier.size = 1) +
  facet_wrap(~animal,ncol=3) + theme_light()
```

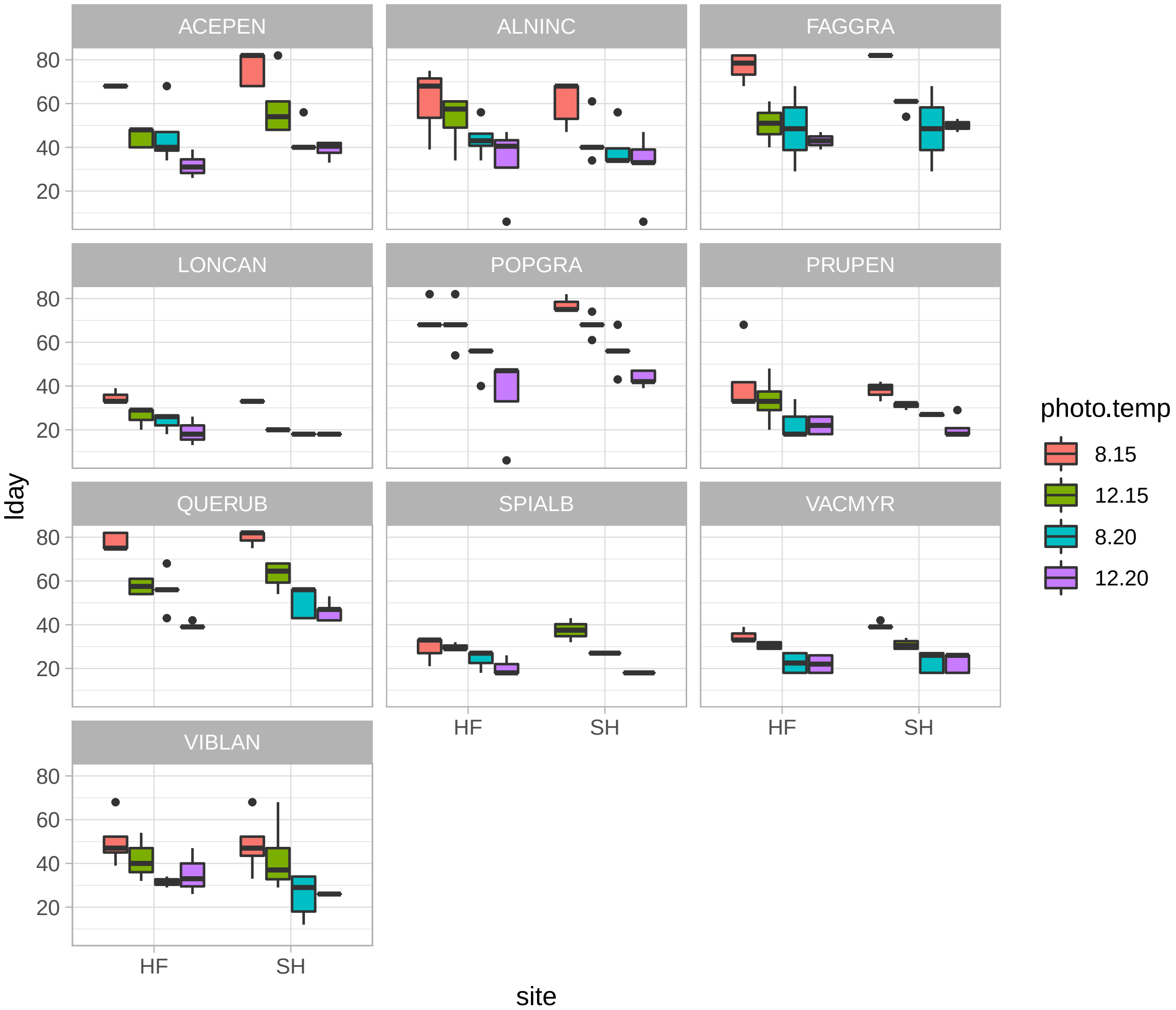

It is interesting to see that in general the photoperiod as a stronger effect at 15 C than at 20 C. This explains the significant interaction. This interaction appears to be slightly species specific, however.

Let's now look at the **random effects**. To evaluate variance explained by the random effects, it is useful to look at the relative proportion of the variance explained by each effect and by the residuals (`units`):

```
# Proportion of variance explained by random factors
rand <- M3$VCV/apply(M3$VCV,1,sum)
# Get median values (50%) and 95% quantiles
apply(rand,2,function(c) quantile(c,probs = c(0.025,0.5,0.975)))
```

```
##          intra.svd.       animal      units
## 2.5%  1.414953e-05 4.209442e-06 0.05343758
## 50%   5.391075e-01 1.533095e-02 0.30459799
## 97.5% 9.326404e-01 7.993061e-01 0.66011431
```

```
# Get the mean value
apply(rand,2,mean)
```

```
## intra.svd.     animal      units
##  0.4783360  0.2092704  0.3123937
```

```r
# Also get 95% quantiles and mean for Heredity (H^2)
rand <- apply(M3$VCV[,1:2],1,sum)/apply(M3$VCV,1,sum)
quantile(rand,probs = c(0.025,0.5,0.975))
```

```
##      2.5%       50%     97.5%
## 0.3398857 0.6954020 0.9465624
```

```r
mean(rand)
```

```
## [1] 0.6876063
```

```r
# And the intraspecific variance relative to the phylogenetic variance
rand <- M3$VCV[,1]/apply(M3$VCV[,1:2],1,sum)
quantile(rand,probs = c(0.025,0.5,0.975))
```

```
##         2.5%          50%        97.5%
## 2.316468e-05 9.760388e-01 9.999946e-01
```

```r
mean(rand)
```

```
## [1] 0.6850599
```

We find that the phylogenetic component of the variance explains ca. 21 % of the variance, that the intraspecific component explains ca. 48%, but that confidence intervals are quite large.

##### Other models

It is interesting to compare these results to models where the intraspecific structure is excluded. Here the model with only the phylogenetic genetic structure:

```r
summary(M1)
```

```
##
##  Iterations = 5001:104981
##  Thinning interval  = 20
##  Sample size  = 5000
##
##  DIC: 2134.531
##
##  G-structure:  ~animal
##
##        post.mean l-95% CI u-95% CI eff.samp
## animal     196.6    48.59    425.7     5000
##
##  R-structure:  ~units
##
##       post.mean l-95% CI u-95% CI eff.samp
## units     88.05    73.89    103.3     5000
##
##  Location effects: lday ~ warm * photo
##
##               post.mean l-95% CI u-95% CI eff.samp  pMCMC
## (Intercept)      55.820   44.740   66.752     5000 <2e-04 ***
## warm20          -19.941  -23.050  -16.983     5000 <2e-04 ***
## photo12         -11.666  -14.782   -8.577     5000 <2e-04 ***
## warm20:photo12    5.360    0.857    9.541     4784 0.0164 *
## ---
```

```
## Signif. codes:  0 '***' 0.001 '**' 0.01 '*' 0.05 '.' 0.1 ' ' 1
```

You can see that the interaction between temperature and photoperiod is similar in terms of effect size (it is only slightly lower), but it is less significant.

And now the model in which no random effects are taken into account.

```
summary(M0)
```

```
##
##  Iterations = 5001:104981
##  Thinning interval  = 20
##  Sample size  = 5000
##
##  DIC: 2425.909
##
##  R-structure:  ~units
##
##       post.mean l-95% CI u-95% CI eff.samp
## units     247.8    207.2      287     5225
##
##  Location effects: lday ~ warm * photo
##
##                 post.mean l-95% CI u-95% CI eff.samp  pMCMC
## (Intercept)       58.104   54.510   61.832     5225 <2e-04 ***
## warm20           -19.679  -24.719  -14.538     5464 <2e-04 ***
## photo12          -10.913  -16.045   -5.644     5000 <2e-04 ***
## warm20:photo12     4.335   -2.789   11.641     5000  0.233
## ---
## Signif. codes:  0 '***' 0.001 '**' 0.01 '*' 0.05 '.' 0.1 ' ' 1
```

Here, the interaction between the temperature and the photoperiod (`warm:photo`) is not significant anymore, and its effect has shrinked!
